# Emergent effects of synaptic connectivity on the dynamics of global and local slow waves in a large-scale thalamocortical network model of the human brain

**DOI:** 10.1101/2023.10.15.562408

**Authors:** Brianna M Marsh, M. Gabriela Navas-Zuloaga, Burke Q Rosen, Yury Sokolov, Jean Erik Delanois, Oscar C González, Giri P. Krishnan, Eric Halgren, Maxim Bazhenov

**Affiliations:** Department of Medicine, University of California, San Diego; Neuroscience Graduate Program, University of California, San Diego; Department of Computer Science and Engineering, University of California, San Diego; Departments of Radiology and Neuroscience, University of California, San Diego

**Author notes:** **For correspondence:** (MB). These authors contributed equally to this work (Brianna M Marsh, M. Gabriela Navas-Zuloaga).

## Abstract

Slow-wave sleep (SWS), characterized by slow oscillations (SO, <1Hz) of alternating active and silent states in the thalamocortical network, is a primary brain state during Non-Rapid Eye Movement (NREM) sleep. In the last two decades, the traditional view of SWS as a global and uniform whole-brain state has been challenged by a growing body of evidence indicating that SO can be local and can coexist with wake-like activity. However, the understanding of how global and local SO emerges from micro-scale neuron dynamics and network connectivity remains unclear. We developed a multi-scale, biophysically realistic human whole-brain thalamocortical network model capable of transitioning between the awake state and slow-wave sleep, and we investigated the role of connectivity in the spatio-temporal dynamics of sleep SO. We found that the overall strength and a relative balance between long and short-range synaptic connections determined the network state. Importantly, for a range of synaptic strengths, the model demonstrated complex mixed SO states, where periods of synchronized global slow-wave activity were intermittent with the periods of asynchronous local slow-waves. Increase of the overall synaptic strength led to synchronized global SO, while decrease of synaptic connectivity produced only local slow-waves that would not propagate beyond local area. These results were compared to human data to validate probable models of biophysically realistic SO. The model producing mixed states provided the best match to the spatial coherence profile and the functional connectivity estimated from human subjects. These findings shed light on how the spatio-temporal properties of SO emerge from local and global cortical connectivity and provide a framework for further exploring the mechanisms and functions of SWS in health and disease.

**Author Summary:** Slow Wave Sleep (SWS) is a primary brain state displayed during Non-Rapid Eye Movement (NREM) sleep. While previously thought of as homogenous waves of activity that sweep across the entire brain, modern research has suggested a more nuanced pattern of activity that can vary between local and global slow wave activity. However, understanding how these states emerge from small scale neuronal dynamics and network connectivity remains unclear. We developed a biophysically realistic model of the human brain capable of generating SWS-like behavior, and investigated the role of connectivity in the spatio-temporal dynamics of these slow waves. We found that the overall strength and a relative balance between long and short-range synaptic connections determined the network behavior - specifically, models with relatively weaker long-range connectivity resulted in mixed states of global and local slow waves. These results were compared to human data, and we found that models producing mixed states provided the best match to the network behavior and functional connectivity of human subject data. These findings shed light on how the spatio-temporal properties of SWS emerge from local and global cortical connectivity and provide a framework for further exploring the mechanisms and functions of SWS in health and disease.

## Introduction

Internal states of the brain rapidly and profoundly influence sensation, cognition, emotion, and action (***Lee and Dan, 2012***; ***Pfaff et al., 2008***; ***Steriade and McCarley, 2013***). Dramatic change of internal brain state occurs at the transition between wakefulness and sleep (***Campbell and Tobler, 1984***; ***Leung and Mourrain, 2018***; ***Steriade and McCarley, 2013***). The main electrophysiological hallmarks of slow wave sleep (SWS) and rapid eye movement (REM) sleep are shared across mammals. During the SWS, the brain dynamics is dominated by oscillations between an active (Up) and a silent (Down) states, the activity called slow oscillation (SO, <1Hz) (***Steriade et al., 1993a***,b). Alterations of these sleep dynamics have significant consequences for brain function and are at the core of many neuropsychiatric disorders (***Winkelman and Lecea, 2020***).

Until recently, sleep/wake transitions or transitions between different stages of sleep were thought to occur uniformly throughout the cortex. Recent evidence in rodents, primates and humans points to the existence of electrographic events resembling local sleep during behavioral wakefulness (***Vyazovskiy et al., 2011***; ***Hung et al., 2013***; ***Quercia et al., 2018***; ***Engel et al., 2016***; ***van Kempen et al., 2021***). During wakefulness, local groups of cortical neurons tend to fall silent for brief periods, as they do during SWS. Similarly, during behavioral sleep, transient local wake-like activity can intrude SWS (***Nobili et al., 2011***; ***Peter-Derex et al., 2015***). SWS and REM sleep can also coexist in different cortical areas (***Soltani et al., 2019***; ***Funk et al., 2016***; ***Bernardi et al., 2018***). Homeostatic sleep pressure can affect cortical activity locally (***Siclari and Tononi, 2017***). Spindles and slow waves, may increase locally after learning (***Eschenko et al., 2006***; ***Pugin et al., 2015***), suggesting experience dependent regulation. Together these results suggest an intriguing idea that local slow-waves may have a functional significance enabling brain to process and learn in a parallel way. However, the underlying mechanisms of local vs global slow-wave dynamics as well as factors controlling transitions between them are unknown.

We previously developed computer models of sleep SO (***Timofeev et al., 2000***; ***Bazhenov et al., 2002***), including transitions between awake and sleep states (***Krishnan et al., 2016***). However, the highly detailed and computationally intensive nature of these models have prevented up-scaling to investigate slow wave dynamics at the whole-brain scale. Recently, several whole-brain models of slow-wave sleep incorporating biologically grounded connectivity from probabilistic diffusion MRI tractography and providing a good fit to human sleep recordings were proposed (***Cakan et al., 2022***; ***Goldman et al., 2023***). These models investigated the role of long-range connections in SO brain-wide propagation. However, the mean-field neural-mass representation of cortical regions in these models provides only limited insights on how local (within a few milimeters) and long-range connectivity and activity interact to affect initiation and propagation of sleep slow waves.

To bridge this gap between micro and macro-scale mechanisms, we have developed a multiscale, whole-brain, thalamocortical network model with biologically grounded cortical connectivity that exhibits the essential activity states of wake and slow wave sleep. The model consists of 10,242 cortical columns per hemishpere, each containing spiking pyramidal (PY) and inhibitory (IN) neurons arrayed in 6 layers, and 642 thalamic cells of each of four types (thalamo-cortical (TC) and reticular (RE), belonging to either core or matrix). To investigate the role of connectivity in emerging SO, we systematically modified the density, range, and relative strength of cortical connections, as well as distance-dependent synaptic delays. The model dynamics were compared to the spatial coherence profiles obtained from human subjects. Our study revealed that a characteristic balance of long and short-range connectivity is necessary to match the data and enable the emergence of complex mixed states of local and global SO (***Nir et al., 2011***; ***Bernardi et al., 2018***; ***Vyazovskiy et al., 2009***). It predicts, that changes in synaptic weights, which likely occur over the course of sleep (***Timofeev and Chauvette, 2017***), may enable transitions between global synchronized and local SO.

## Results

### Connectivity

We start with a brief overview of the model (Figure 1, see Methods for details). The model includes 10,242 cortical columns positioned uniformly across the surface of one hemisphere, corresponding to the vertices in the ico5 cortical reconstruction reported in (***Rosen et al., 2019***). The medial wall includes 870 of these vertices, so all analyses for this one-hemisphere model were done on the remaining 9372 columns. Each column has 6 cortical layers (L2, L3, L4, L5a, L5b and L6), with 1 pyramidal cell and 1 inhibitory cell per layer. Cortical neurons are modeled using computationally efficient map-based (***Komarov et al., 2018***) spiking PY and IN neurons. Intra-columnar connections follow the canonical cortical circuit (Figure 1A), while long-range cortical connectivity (Figure 1B) is based on diffusion MRI (dMRI) tractography (***Rosen and Halgren, 2021***) of the Human Connectome Project (HPC) young adult dataset (***Van Essen et al., 2013***), organized into the 180 parcels of the HCP multimodal atlas (***Glasser et al., 2016***)). The originating and terminating layers of long range connections are assigned according to the parcel’s relative hierarchical positions (Figure 1D-E). Column-wise connectivity is obtained by applying the parcel-wise relation between connection length and probability to synthesized intercolumnar distances (Figure 1C, see Diffusion MRI Guided Connection Probability for details). The resulting connectivity (Figure 1G) retains the parcel-wise structure from the data (***Rosen and Halgren, 2021***) (Figure 1F, Pearson’s correlation r = 0.81, p < 0.0001), with an approximately exponential distribution between connection frequency and length (see Figure 1C). The connection delays, which are proportional to the geodesic connection lengths, follow a similar distribution. The thalamus is simulated with layers of thalamocortical (TC) and reticular thalamic (RE) neurons (***Bazhenov et al., 1998a***,b, ***2002***), and consists of the core and the matrix (***Bonjean et al., 2011***, ***2012***), which selectively project to the neurons from different cortical layers satisfying the outlined cortico-thalamo-cortical loop (***Shepherd and Yamawaki, 2021***). Following (***Krishnan et al., 2016***) the model included the effects of neuromodulators to induce transitions between Wake state and SWS.

**Figure 1.**
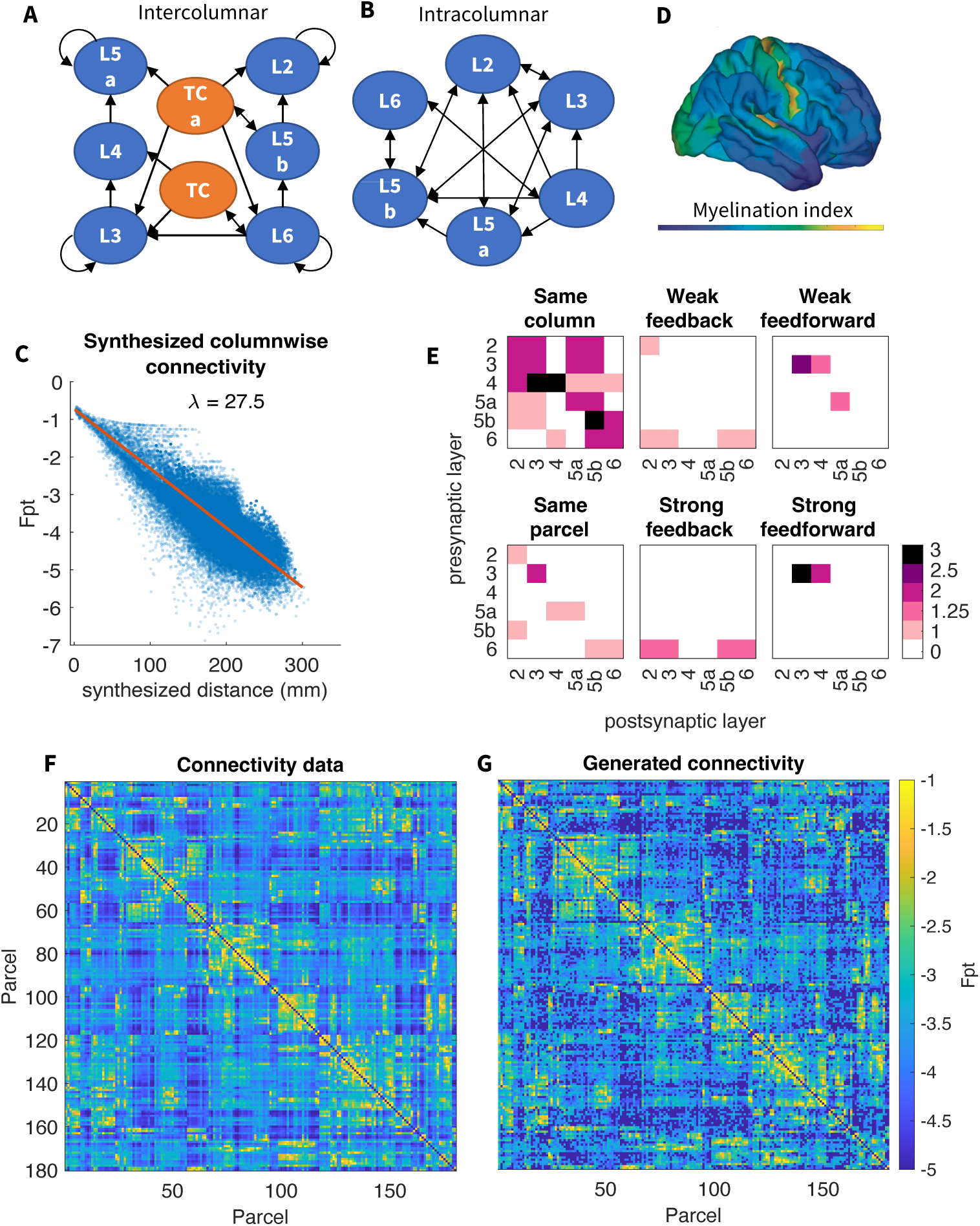
Model architecture. A) Connectivity between neurons in the same column follows the canonical circuit. Arrows indicate directed connections between layers. Neurons are never connected to themselves. B) Connectivity diagram between different cortical columns and thalamocortical cells in the core (TC) or matrix (TCa). C) Exponential decay of connection probability with distance between columns. Previously reported inter-parcel dMRI connectivity (***Rosen and Halgren,*** 2021) shows an exponential relation with fiber distance. Column-wise connection probability is obtained by applying the dMRI exponential function to inter-column fiber distances (synthesized from their corresponding geodesic distances) and then adding the residual inter-parcel dMRI connectivity, estimated by regressing out the exponential trend (see Diffusion MRI Guided Connection Probability for details). D) Estimated myelination index (***Glasser and Essen,*** 2011; ***Burt et al.,*** 2018) throughout the cortex. A hierarchical index inversely proportional to this myelination index is assigned to each of the 180 cortical parcels designated in the HCP-MMP atlas (***Glasser et al.,*** 2016). E) Excitatory corticocortical connections belong to one of 6 classes based on the hierarchical index of the pre- and post-synaptic neurons: within the same column, within the same parcel, weakly or strongly feedforward (from lower to higher hierarchical index) and weakly or strongly feedback (from higher to lower hierarchical index). Based on experimental reports on connectivity (***Markov et al.,*** 2013; ***Rockland,*** 2019) weights are scaled according to the connection type by the factor shown in the matrices. F) Parcel-wise connectivity from ***Rosen and Halgren*** (2021). G) Generated model connectivity, which retains the parcel-wise structure from data in panel F (Pearson’s correlation r = 0.81, p < 0.0001).

In the following sections, we first discuss "baseline" awake and sleep state dynamics of the model. Next we discuss how sleep dynamics are affected by altering network connectivity and reveal conditions for coexistence of local and global SO. We finally present analysis of human recordings and compare different model dynamics to human data using coherence analysis across frequency and distance and functional connectivity analysis. In addition to the basic characteristics of SO frequency, amplitude, and propagation speed, network effects are primarily described in terms of 3 key metrics: *synchrony*, *spread*, and *participation*. Synchrony refers to the timing of the Up state onsets and offsets across all neurons participating in a slow wave. Spread refers to how far the slow wave travels across the cortex. Finally, participation refers to the percent of neurons that participate in a slow wave within the area of spread.

### Wake State Characterization

Wake state (Figure 2A) was achieved by applying biologically relevant changes in key parameter values (Table 4), based on our previous work (***Krishnan et al., 2016***). The resultant network shows a consistent average firing rate of individual neurons at about 17 Hz. Given the broad distribution of firing rates reported in awake state (***Miyawaki et al., 2019***; ***Thomas et al., 2020***), we believe this model is appropriate. There is indeed experimental evidence of average firing rates in awake state being 15 Hz (***Vyazovskiy et al., 2009***), 20-50 Hz (***Thomas et al., 2020***), or 5-16 Hz (***McKillop et al., 2018***). However, it should be noted that different mean firing rates can be achieved with minimal adjustments to key parameters, namely AMPA and GABA strength. The power spectral density (Figure 2a.5) revealed a distinct 1∕*f* phenomenon, as is common for in vivo recordings in awake state (***Donoghue et al., 2020***). Individual neurons showed irregular tonic firing (Figure 2a.1), with baseline membrane voltage around -60 mV (Figure 2a.4). The firing activity was not synchronized (Figure 2a.3), which was reflected in the low voltage LFP (Figure 2a.2). Overall, the network revealed properties representative of biological awake state activity.

**Figure 2.**
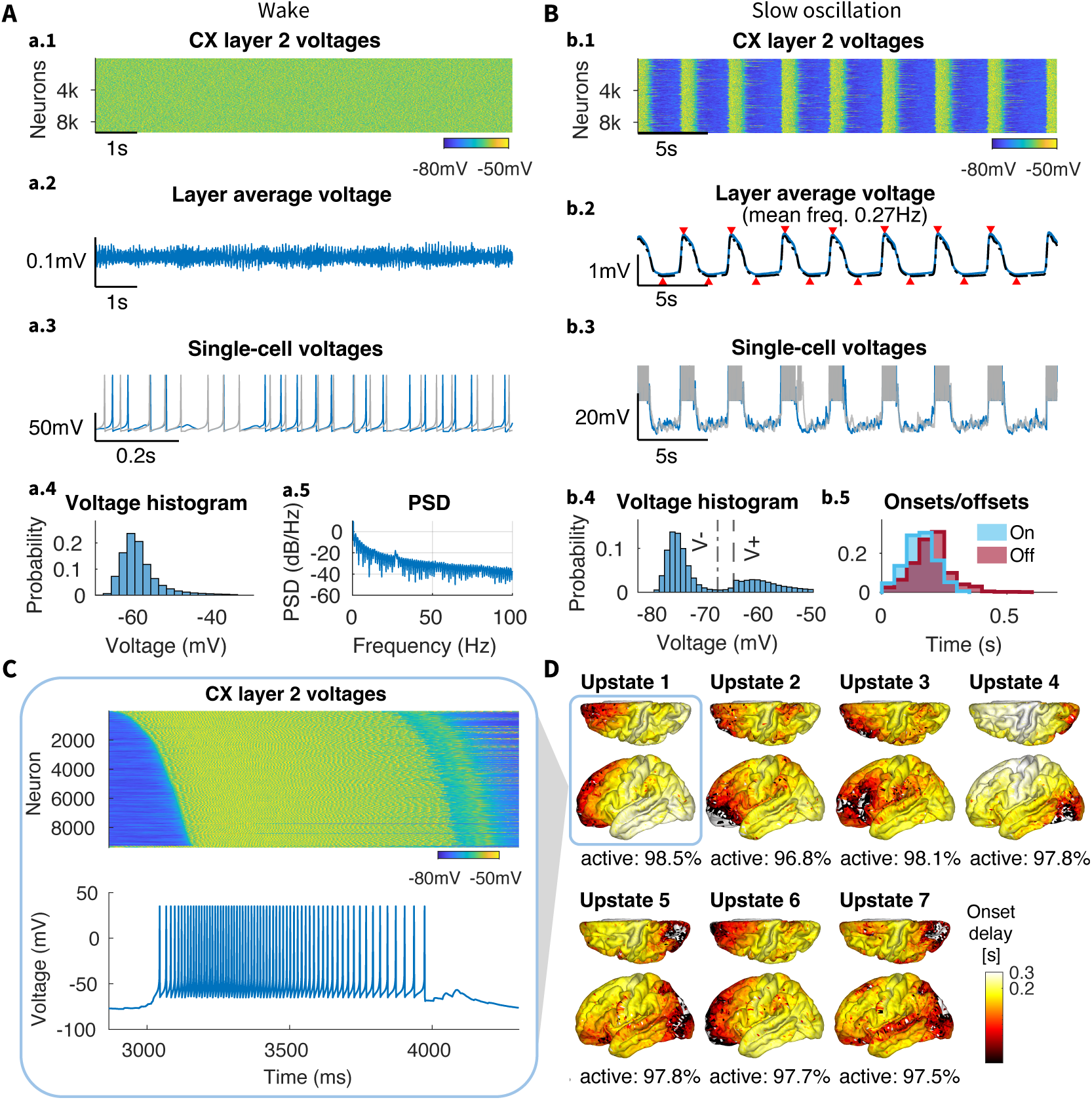
Baseline model activity in awake and sleep states. A) Wake state. a.1) Membrane voltages of all neurons from layer 2 over 5 seconds of simulation; a.2) Layer average voltage over time, simulating LFP; a.3) Activity of two representative neurons from layer 2, both showing irregular tonic firing; a.4) Voltage histogram of all neurons over the whole simulation time, note approximately -60mV peak representing baseline membrane voltage; a.5) Power spectral density (PSD) of the average voltage revealed a distinct 1∕*f* phenomenon typical for *in-vivo* recordings. B) Slow-wave dynamics. b.1) Membrane voltages of all neurons (excluding medial wall) from layer 2 over 30 seconds of simulation, revealed synchronized bands of activity during Up states; b.2) Average voltage of all cortical layers (*dashed black line*) and layer 2 neurons (*solid blue line*, excluding medial wall) over time, with nearly identical behavior. *Red triangles* above and below the trace mark global Up and Down states, respectively, from layer 2 (coincident with global Up and Down states from the average of layers); a.3) Activity of two representative neurons from layer 2, both showing synchronized transitions between Up and Down states; a.4) Voltage histogram of all neurons over the whole simulation time, revealed the characteristic bi-modal distribution caused by Up and Down state alternations during SWS. Dashed vertical lines labeled *V* ^−^ and *V* ^+^ indicate the voltage thresholds used to detect Down states and Up states, respectively; b.5) Distribution of the Up state onsets and offsets for all neurons over the whole simulation time. Narrow histograms indicate highly synchronous initiation and termination of the Up states. C) Zoom into the first Up state from panel b.1, with neurons sorted from earliest to latest onset time, and a single cell voltage from panel b.2, showing the transition from Down to Up state, steady firing during the Up state, and transition to Down state. D) Latency map, calculated as the onset delay of each neuron with respect to the earliest onset, for each Up state in panel b.2 (see Onset/offset Detection in Methods for details). The percent of active neurons during each Up state is shown below the corresponding latency map. Up states involve nearly every neuron in the cortex within 300ms from its initiation time.

### Slow Wave Characterization

To attain a network displaying SWS-like activity from the wake state baseline, several cellular and neurochemical parameters were adjusted according to the methodology presented in (***Krishnan et al., 2016***). Namely, we increased cortical AMPA and GABA conductances as well as the leak current and slow hyperpolarizing currents in all cortical neurons (see Methods, ***Table 4***). The resulting baseline model of SWS (Figure 2B) revealed synchronized and regular oscillations (∼0.27 Hz) between silent (Down) and active (Up) states (Figure 2b.2) in all cortical excitatory and inhibitory neurons (Figure 2b.1). This network dynamics was organized as the waves of spiking activity propagating across the entire cortex - slow-waves. (This can be seen in Supplemental Video S1.) The slow-waves were further highly synchronized across layers (Supplementary Figure S1). Differences between layers include a relatively less active L4. This is probably due to L4 receiving within-column connections from only L6, while other layers have inputs from two or more other layers (see Figure 1E).

In general, the network spent more time in the Down states, and the Up state onsets and offsets were well synchronized (see narrow onset and offset histograms in Figure 2b.5). This pattern of longer Down states is typical for recordings under urethane anesthesia (***Steriade et al., 1993a***), while Down states may be shorter in non-anesthetized animals (***Steriade et al., 2001***). Nevertheless, the frequency of SOs was within the range of human data (***Csercsa et al., 2010***). The latency maps in Figure 2D demonstrate how waves of activity travel from a single initiation site, which varied between Up states with no detectable dependency on the previous ones. Each Up state involved >97% of all excitatory neurons. In this baseline model, slow waves revealed both high spread and high participation.

Cortical inhibitory, thalamocortical, and reticular thalamic neurons all showed similar patterns of oscillations (Supplementary Figure S2), as previously shown in ***Bazhenov et al.*** (***2002***). While thalamic cells followed cortical oscillations, they were not directly involved in the generation of SO. In fact, simulations with no thalamic inputs to the cortex maintained cortical SO (Supplementary Figure S5). This is in line with experimental results showing that SO survives extensive thalamic lesions (***Steriade et al., 1993a***), and is present in neocortical slices (***Sanchez-Vives and McCormick, 2000***) and isolated cortical slabs (***Timofeev et al., 2000***). Nevertheless, removing the thalamus reduced synchrony of cortical slow waves (see Supplementary Information: Effect of thalamus on SO synchronization), in agreement with in vivo data (***Lemieux et al., 2014***).

### Connection Density

In the next two sections, we test effects of alternating the intracortical connection density and the maximal radius of connections on SO properties. Importantly, in these simulations we scaled the strengths of remaining synaptic connections to compensate for the lost connections and to ensure an equivalent amount of net activation per neuron (see Modifying Network Structure for details).

In the baseline model, synaptic connectivity was set to match biological data (see Methods). Next, to test the effects of connection density between regions, we gradually changed the density of connections, by applying a probability *P* to retain each possible connection (Figure 3A-B) regardless of the distance between neurons, thus keeping the connection distance distribution unchanged (Figure 3C). Remaining synapses were scaled to keep total synaptic input per neuron unchanged.

**Figure 3.**
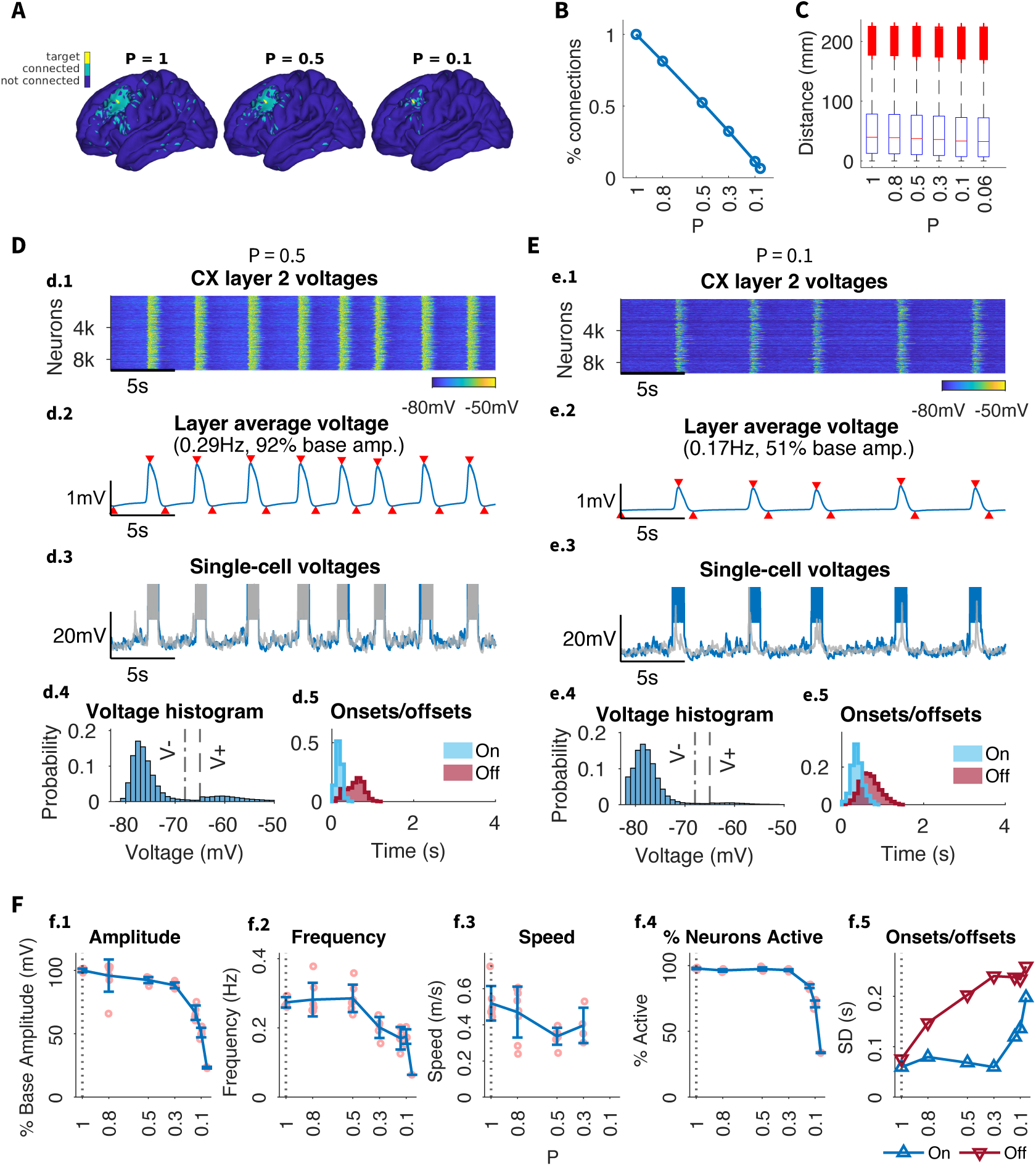
Effect of connection density on SO dynamics. A) Example target cortical column ( *yellow*) with all columns connected to it through any layer (*green*) for different connection densities. Density is reduced by decreasing *P*, which denotes the probability of preserving a connection from the original dMRI-based connectivity. With *P* = 1 all connections are preserved, while for *P* = 0.5, *P* = 0.3 and *P* = 0.1 each connection is preserved with a 50%, 30% or 10% probability, respectively. B) Percent of the original number of connections retained for different values of *P*. C) Distribution of connection distances, or lengths, for different values of *P*. Red horizontal lines indicate medians, bottom and top box edges indicate the 25th and 75th percentiles, respectively, whiskers extend to the most extreme data points not considered outliers, and outliers are plotted individually using the ‘+’ marker symbol. Note, connection density is reduced uniformly across all lengths. D-E) Individual analysis of SO dynamics for different values of *P*. Subpanels as in Figure 2. F) Summary of the effect of reducing connection density on the global SO frequency and amplitude as well as the standard deviation of the onset/offset delays (i.e. the width of the onset/offset histograms in d.5 / e.5)

Setting the probability from 100% down to 50% (i.e., ablating a random half of all connections) had minimal effect on the slow waves, indicating the model’s robustness to even significant loss of individual connections as long as the total strength of excitatory synaptic input per neuron remains unchanged. Slow waves exhibited a similar propagation pattern with only a slightly reduced amplitude (approx. 92% of baseline SOs) and speed (Figure 3D, F). The synchrony of the Up state offsets, however, was impaired; the distribution of offset timings across neurons became much wider (Figure 3d.5, f.5) compared to baseline model (Figure 2b.5). Further decreasing density below 30% caused a considerable drop in amplitude and participation (Figure 3f.1, f.4). In the extreme case of only 10% of connections remaining (*P* = 0.1, Figure 3E) (see Supplementary Information: Effects of connections density, range and delays and Figure S6), there was continued loss of synchrony in Up state offsets and onsets, slow waves became less frequent (down to 0.17 Hz), with lower amplitude (50% of baseline), and lower participation (only 70% of neurons involved per Up state) (Figure 3F). Nevertheless, slow waves still propagated through the all brain regions on the macro scale, showing that long range synchrony is still possible (if slightly impaired) with only very few connections in place as long as those connections are sufficiently strong and long-range connections are present. Below this level of remaining connections, slow waves no longer occurred and the network was largely silent.

### Connection Range

To determine the specific contribution of long-range connections, we varied the maximum connection radius; any connections longer than a specified distance *R* were set to 0 (Figure 4A). Again, the strengths of remaining connections were scaled up to compensate for reduction in the total number of connections and to keep total synaptic input per cell unchanged (see Modifying Network Structure for details). In contrast to the density reduction (see Figure 3C), this alteration did shift the connection distance distribution (Figure 4C).

**Figure 4.**
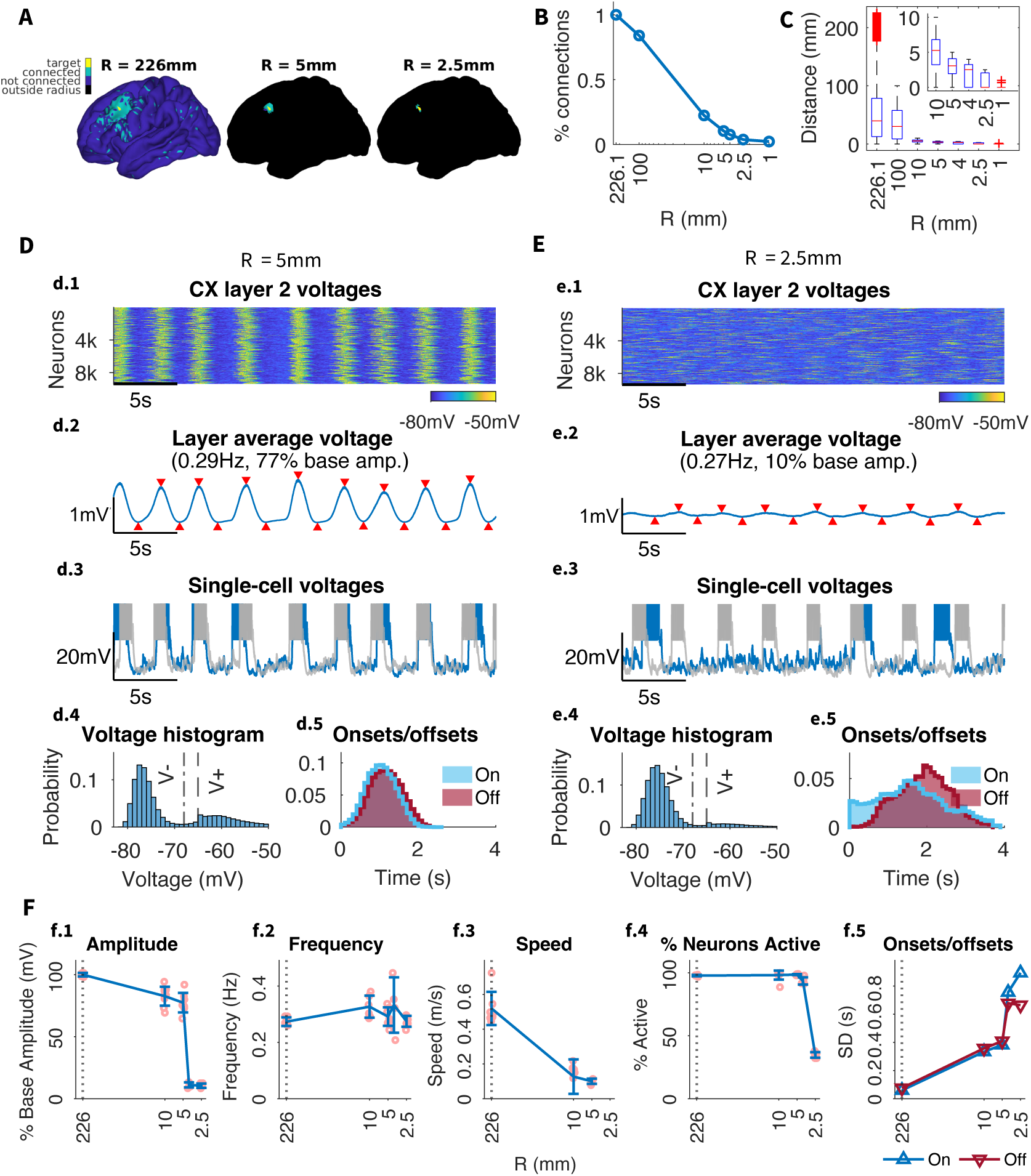
Effect of connection range on SO dynamics. A) Example target column (*yellow*) with all its connected columns (*green*). The *blue* area indicates the connection range for the corresponding radius *R*. No connections are made in the *black* region outside the radius. *R* = 226*mm* encompasses all the original dMRI-derived connections. B) Percent of the original number of connections preserved for different values of *R*. C) Distance distribution for different values of *R*, with *inset* zooming into radii below 10mm. *R* imposes a maximum connection length and truncates the distance distribution accordingly. Red horizontal lines indicate medians, bottom and top box edges indicate the 25th and 75th percentiles, respectively, whiskers extend to the most extreme data points not considered outliers, and outliers are plotted individually using the ‘+’ marker symbol. D-E) SO analysis for *R* = 5*mm* and *R* = 2.5*mm*. Subpanels as in Figure 2. F) Summary of the effect of reducing connection density on the global SO frequency and amplitude as well as the standard deviation of the onset/offset delays (i.e. the width of the onset/offset histograms in d.5 / e.5)

The maximum connection length in baseline model was 226.1 mm, but most connections (approximately 80%) were maintained within 100 mm (Figure 4B). A connection radius as small as 5 mm, which includes second-order neighbors of a given neuron, resulted in only approximately 10% of connections still intact. The alterations of the maximum connection length led to significantly different activity patterns than in the network with a spatially uniform drop to the same percentage of connections (Connection Density). The network with a maximum connection radius of 5 mm (Figure 4D) showed no reduction in the frequency of the slow waves, but both the onsets and offsets of the slow waves revealed dramatic reduction in synchrony. Additionally, there was a substantial drop in slow wave propagation speed, while changes in connection density had minimal effect on speed (compare Figure 4F and Figure 3F). Despite all this, characteristic properties of global slow waves (high spread and high participation) found in the baseline model were maintained.

However, reducing radius below 5mm (i.e., not including 2nd degree neighbors) caused a dramatic drop in global oscillation amplitude, akin to a phase transition from global to local oscillations. With a connection radius of 2.5 mm (Figure 4E), including only first order neighbors of each cell, global slow waves were lost in favor of local slow waves (Figure 5, low spread, high participation). Note that this radius retains only 3.7% of connections, where a spatially uniform drop to any less than 10% of connections removed slow waves entirely. The local slow waves were generated independently in few selected areas, and occasionally showed regional propagation but never travelled across the full hemisphere (Figure 5F). This resulted in reduced global SO amplitude as longrange synchrony between locally oscillating neural populations was impaired (Figure 5C-D). This model can further be seen in Supplemental Video S3. While a large fraction of the cortex was not participating in SO, the active regions (such as region 6, Figure 5) were consistent across Up states. A connectivity analysis revealed that these regions correspond to the network areas with the highest node degree (see Supplementary Figure S9). The same regions exhibited local slow waves for different simulations with the same connectivity parameters but different random seeds for network generation, implying that these regions are not random but emerge from the underlying structure imposed by the dMRI connectivity data.

**Figure 5.**
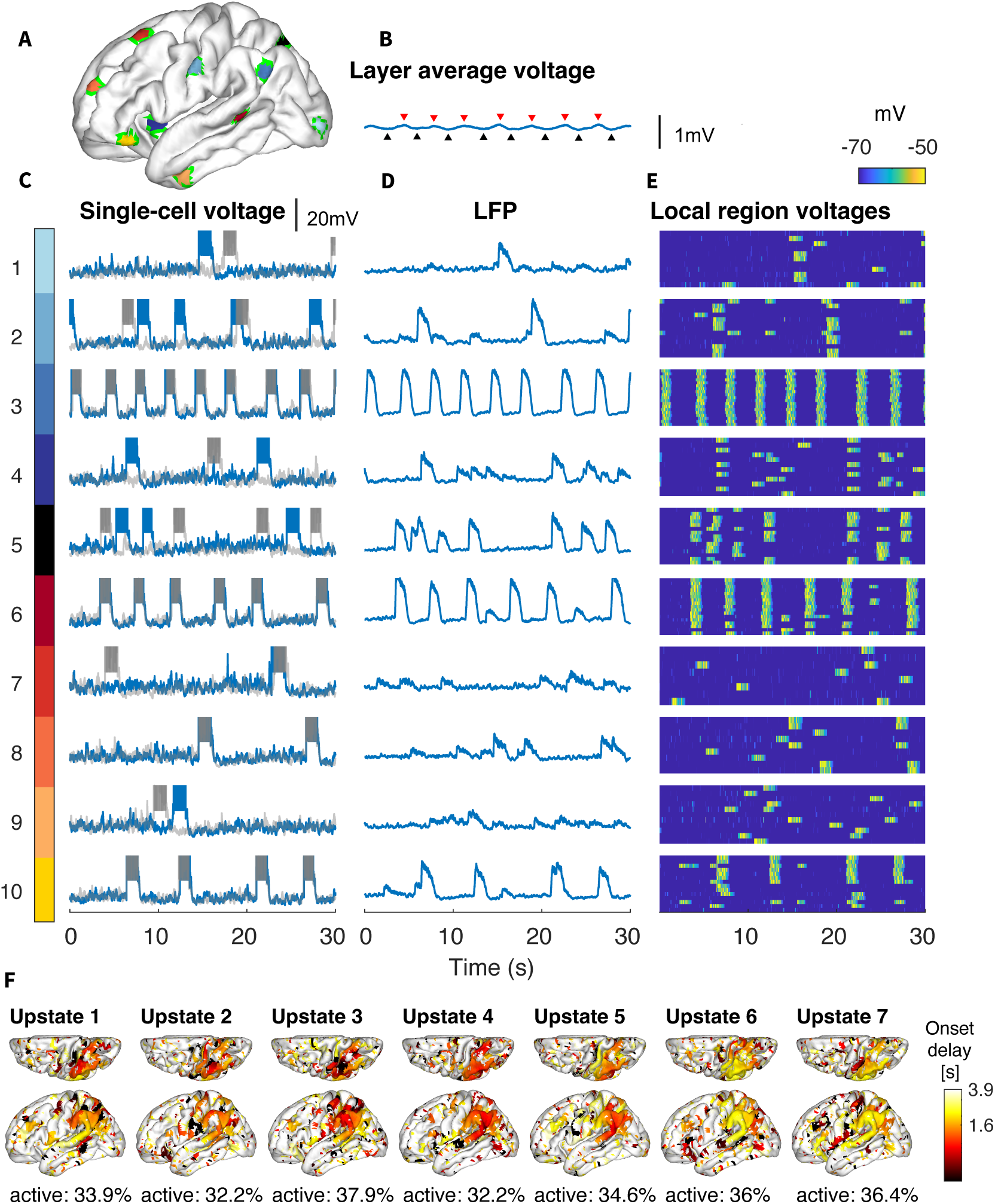
Local SO for small connection radius (*R* = 2.5mm). A) Ten cortical areas with a 5mm radius, that were used to calculate local dynamics. B) Average membrane voltage of layer II neurons, as in Figure 4e.2. C-E) For each region in (A), subpanels show (C) the single-cell voltage for two neurons in the area, (D) the local field potential (LFP) for the 5mm area, and (E) heatmap of individual voltages of all neurons in the area. Note that even within these very small regions, there are still subgroups of independently synchronized neurons. F) Latency map, calculated as the onset delay of each neuron with respect to the earliest onset, for each Up state identified in Figure 4e.2. Note that even the most global of slow waves has very low participation, around 35%, and can be seen to consist of many extremely small initiation sites that generally do not spread.

As a whole, this analysis revealed that long range connections (longer than 5 mm) are essential for the large-scale synchrony of the slow waves, but having only local connections (less than 5 mm) is still sufficient to make slow waves propagating over entire cortex (as long as remaining connections strength is scaled up to maintain total synaptic input per cell). Even with only first order neighbors, the network still maintained slow wave initiation, synchrony, and participation with local propagation.

### Connection Delay

The model implements synaptic delays scaled by geodesic distance between neurons with a maximum delay of 40 ms. The delay for the majority of synapses is under 2 ms because the vast majority of connections are relatively short range. To test the effects of delay times, we changed the delays in two ways (Figure S7): 1) by changing the maximum delay and scaling all others proportionally, and 2) by setting one uniform delay to all connections.

Setting the maximum delay to be smaller (all the way down to 0.1 ms) had little to no effect on slow wave amplitude, frequency, or synchrony. Setting the maximum delay to be larger also had minimal effect on network activity until biologically implausible delays were implemented (≥1 second). Even at this extreme the network retained strong global slow waves, albeit with slightly lower amplitude, longer frequency, and de-synchronized onsets and offsets. Setting a uniform delay of 10 ms (where most delays were previously under 2 ms) further left the network behavior largely unchanged. Only when the uniform delay was increased to 30 ms did we observe a loss of amplitude and synchrony (Figure S7). Since the delay times here shown to have an effect on network behavior are clearly beyond the range of biological plausibility, these results indicate that synaptic delays are not a primary determinant of slow wave spatio-temporal properties.

### Connection Strength

To probe the effects of absolute connection strength on the network activity, we varied the base connection strength between all cortical pyramidal neurons (Figure 6A, C). In contrast to experiments with changing connectivity density or radius, where remaining synapses were scaled to keep total synaptic input constant, this changed the total synaptic input per cell. Weight reductions or increments by a factor of 4 or more caused the slow oscillations to be barely detectable or undetectable (note the low amplitude at the extremes of Figure 6c.1). This was a result of low activity in the case of weights reduction, and of sustained Up state (Down state absence) in the case of weights increase. In general, within the detectable range, the SO amplitude, frequency, and participation decreased when synaptic strength was decreased (Figure 6C). Note that the model revealed an overall robustness to weight variation: the slow-wave propagation was minimally altered by strength increases or decreases by a factor of 2.

**Figure 6.**
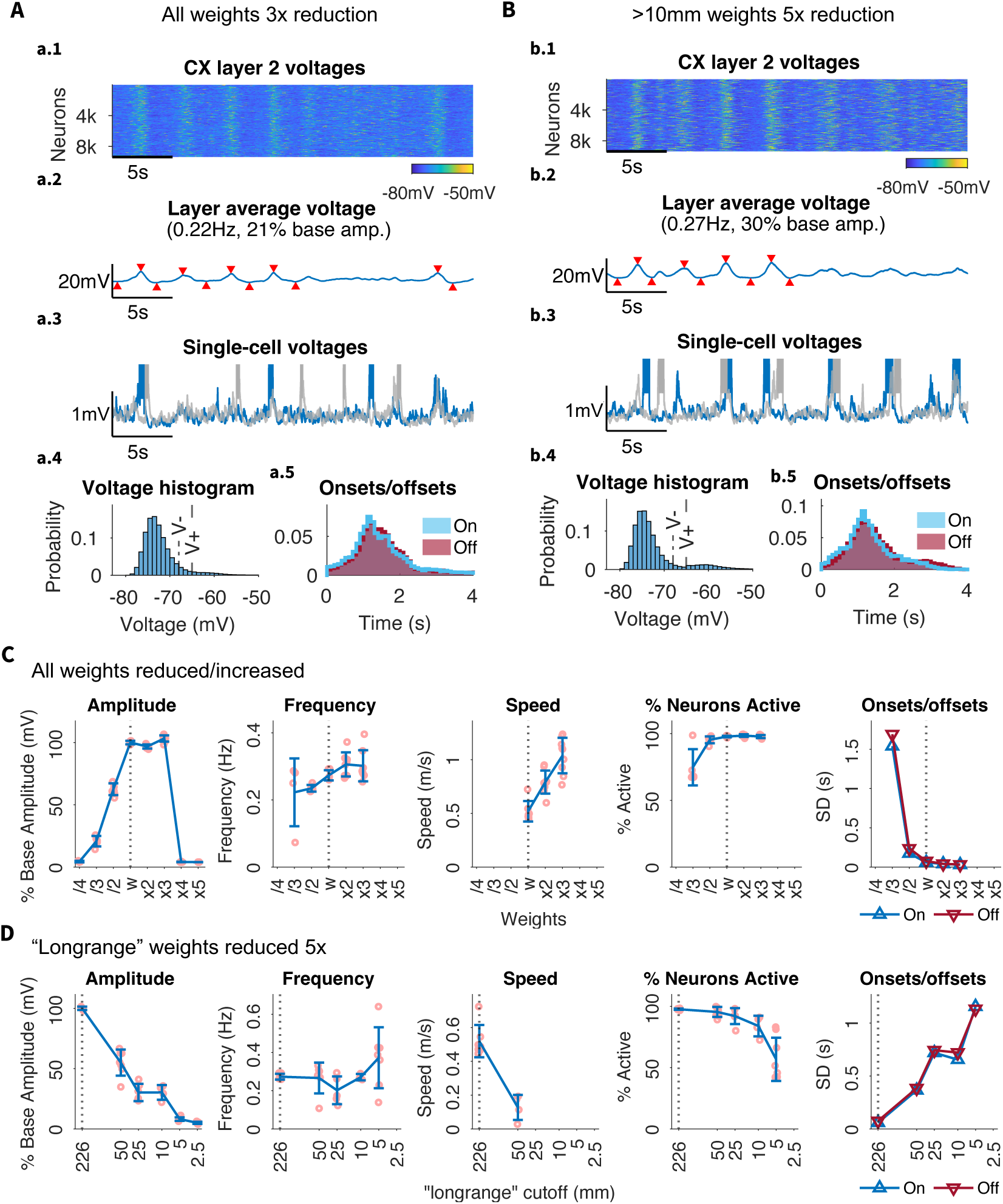
Effect of cortical excitatory synaptic strength on the network dynamics. A) Reduction of all cortical connections by 3x. B) Reduction of only connections longer than 10 mm by 5x. All subpanels as in Figure 2. C) Summary of changes in frequency, amplitude, speed, participation, and onset/offset distribution spread when all connections are reduced as shown in (A). Increasing all weights shows little effect while decreasing all weights shows decreased amplitude and participation, with increases in onset/offset distributions (decreases in synchrony). The frequency of slow waves also becomes more variable. Note, Increasing all weights 5x results in the ablation of slow waves due to constant Up state, i.e., SO characteristics cannot be meaningfully quantified. D) Summary of changes in SO characteristics when only long range connections are reduced as shown in (B). Across all plots, 5 mm is seen to be an inflection point (or "elbow") where network activity changes.

To further investigate the role of long range connectivity, we next performed *N* times synaptic weight reductions on only "long-range connections", i.e., connections over a defined minimum radius *R*_*th*_ (Figure 6B,D). We varied the scaling factor *N* from 2 to 7 and radius threshold *R*_*th*_ from 2.5 mm to 50 mm. (Figure 6B illustrates the case *N*=5, *R*_*th*_=10mm). Again, we did not scale the total synaptic input per cell, so these alternations affected balance of excitation and inhibition in the network. This experiment can be interpreted to model the effect of stronger synaptic inputs nearest to the cell body (***Hawkins and Ahmad, 2016***; ***London and Segev, 2001***).

When connections over *R*_*th*_=10mm were significantly weakened (*N*=5) (Figure 6B, Figure 7), the model displayed a number of clearly identifiable local slow waves originating and reliably traveling around the hemisphere in different directions (Figure 7F). Occasionally, these local slow waves would coincide to produce a global slow wave in the LFP ( see Figure 6b.2, where red downward triangles mark the four detectable global UP states analyzed in Figure 7F). In this model, the local populations of neurons would regularly synchronize producing local slow waves but the global synchrony was lost in many cases except for a subset of all events when global slow waves were generated (Figure 7B, E). This was further confirmed by running longer 150 sec simulation (see Supplemental Video S4) that revealed multiple transitions between local and global synchrony states. With reduced spread and high participation, the local slow wave behavior resembles the results from simulations with exclusively local connectivity (see Connection Range), but stems from the more biologically plausible assumption of weaker (rather than nonexistent) long range connectivity. Indeed, the propagation patterns in Figure 7 is similar to the proposed pattern of biological sleep including a mix of Type I (global) and Type II (local) slow waves, as suggested by (***Siclari et al., 2014***; ***Nir et al., 2011***; ***Bernardi et al., 2018***; ***Funk et al., 2016***; ***Vyazovskiy et al., 2007***, ***2011***; ***Riedner*** *et al., 2007*).

**Figure 7.**
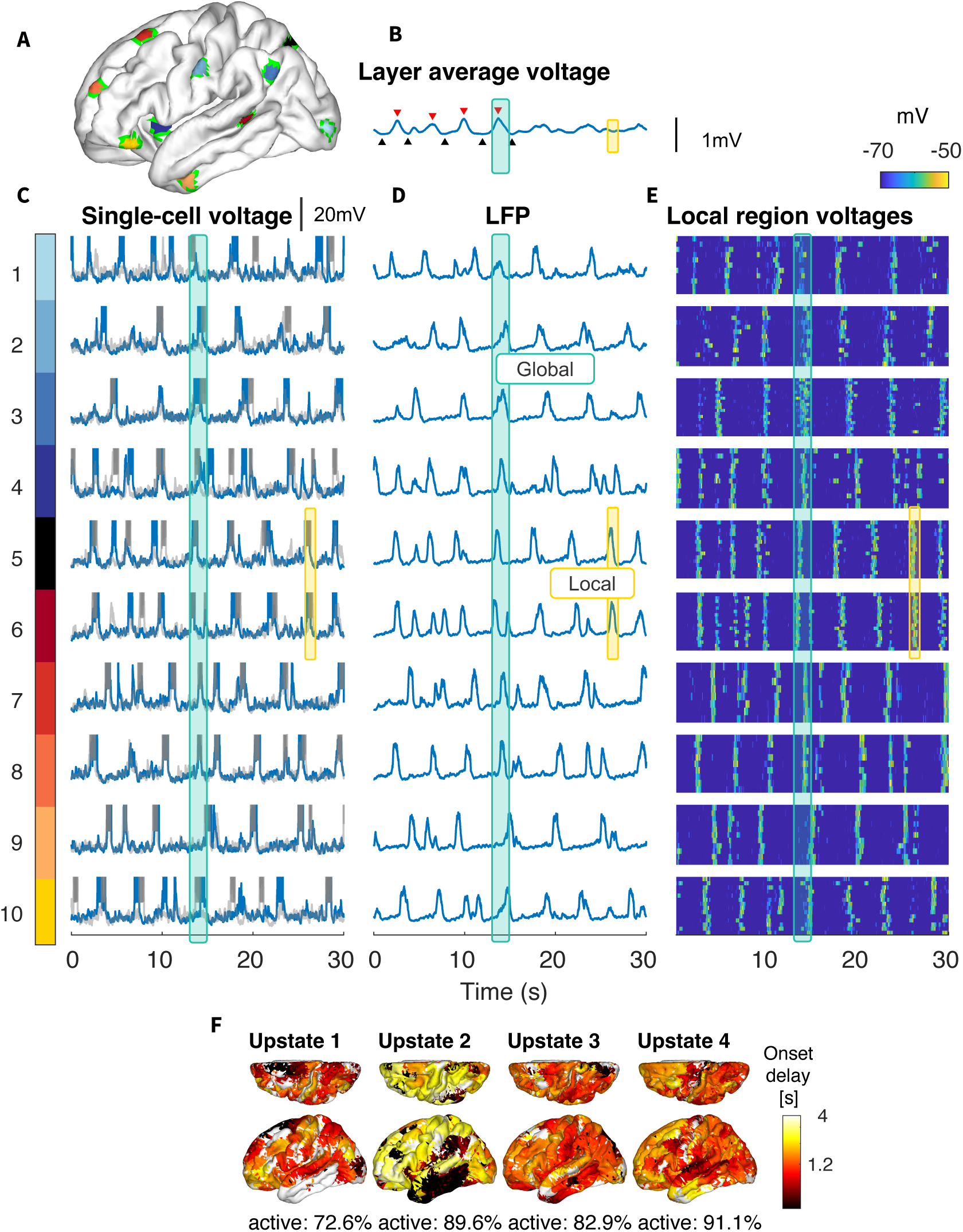
Mixed global (Type I) and local (Type II) Slow Waves: Connections greater than 10mm reduced 5-fold. A) Ten cortical areas with a 5mm radius, that were used to calculate local dynamics. B) Average membrane voltage of layer II neurons. C-E) For each region in (A), subpanels show (C) the single-cell voltage for two neurons in the area, (D) the local field potential (LFP) for the 5mm area, and (E) heatmap of individual voltages of all neurons in the area. This demonstrates that while there is some alignment, up states are not strongly global. F) Latency map, calculated as the onset delay of each neuron with respect to the earliest onset, for each Up state. Note, that while some Up states are global (characteristic examples are marked by blue lines), others are comprised of several independent but coinciding local events (examples are marked by yellow lines).

Importantly, not all the cortical regions were equally likely to generate local slow waves. As in simulations with only local connections, the network areas with the highest node degree (see Supplementary Figure S9) produced almost regular local slow-waves (e.g., region 6, (Figure 7)), while some other (e.g., region 9, (Figure 7)) displayed low frequency irregular spiking with only rare Up state-like events. This again suggests that the active, i.e., generating local SO, areas emerge from the underlying structure imposed by the dMRI-based connectivity.

### Local vs Global Slow-Waves

Given above analysis, we next sought to quantify the global v.s. local slow waves in the model as a function of synaptic strength. For each model, we calculated the regularity in each of the 180 parcels, where regularity is defined as the inverse of approximate entropy. The mean and variance of this distribution are then plotted against each other, as shown in Figure 8A. We define global models as models with high regularity, local models as those with low regularity, and mixed states falling in between. The Intact Score of each model, defined as the percentage of remaining synaptic strength compared to the base model, is further predictive of the regularity and network behavior (Figure 8B). Two groups of models in the middle of distribution with Intact Score 30-50% and regularity 4-12, were identified as mixed states models. In Figure 8C, ten random parcel LFPs from representative global, mixed, and local models show demonstrably different behavior - the global model parcels are all in synchrony, while the local model parcels can be seen to transit into Up states largely independently. The mixed model shows a combination of these behaviors, i.e., transitions to Up state sometimes occurred independently and sometimes in sync across parcels. This group predominately includes models with long-range connectivity scaled down beyond certain radius.

**Figure 8.**
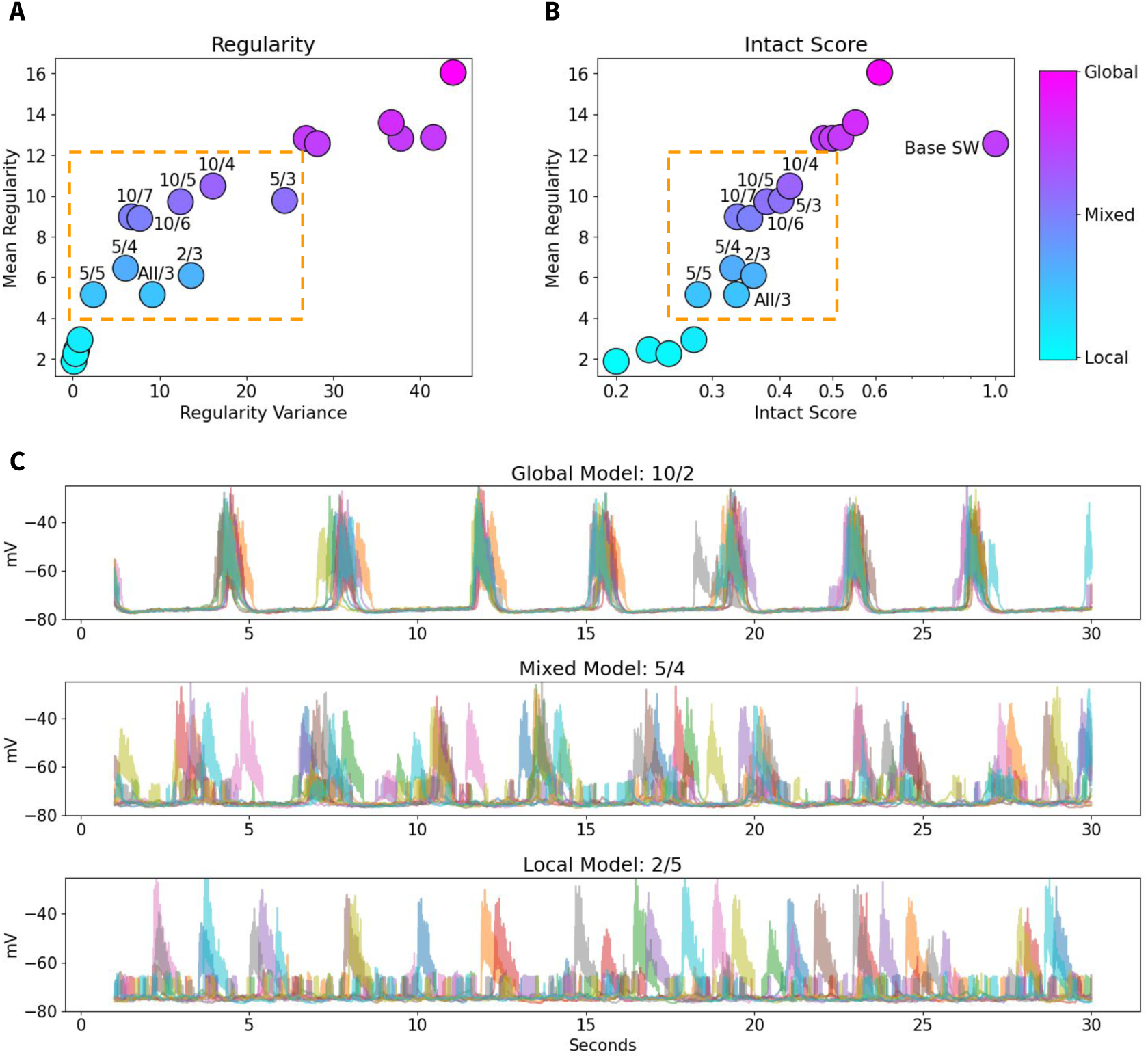
Quantification of Global vs Local Slow Waves. The labels show the radius and factor of strength reduction in each model - e.g., 5/4 indicates a model where connections longer than 5 mm are reduced by a factor of 4. A) For each model, the Regularity of each parcel is calculated; the mean and variance of these distributions predict the overall network behavior, with global models showing high Regularity and local models showing low Regularity. The group of models in the middle of the distribution, boxed in orange, represents Mixed states. B) Mean Regularity is plotted against the Intact Score (total percent of synaptic strength relative to the global base model). Note that the total level of synaptic strength is itself predictive of network behavior - the same mixed models fall in the orange box in the middle of the distribution. C) Example traces of 10 random parcels in representative Global, Mixed, and Local models show different levels of synchrony across the brain.

### Coherence Analysis

To validate our model, we conducted a comparative analysis with experimental data. Specifically, we analyzed coherence across 180 cortical parcels. We calculated coherence in successive narrow-band signals across all parcel pairs to generate a snapshot of coherence across both frequency and distance. We then took the average of the coherence matrices in the SO range (<2 Hz), scaled the values by parcel-to-parcel distances, and masked the model data to match the sparsity of the experimental data. These modified 2D coherence matrices were then used to calculate 2 metrics of functional connectivity: Percent connected and number of communities. Percent connected reports the percentage of parcel pairs with a coherence greater than a given threshold, while the number of communities is determined using Louvain community detection.

Subsequently, we fit the full coherence matrices (across all distances and frequencies) with an exponential model. From this model, we estimated the spatial length constant (*λ*) and the scaling term (*α*). These four metrics were used to compare the functional connectivity and coherence profiles of different biophysical models against in vivo data. Full details of coherence analysis can be found in the Methods under Coherence Analysis.

#### Empirical coherence

The experimental data collection is fully described in Experimental Data. Briefly, continuous stereo-electroencephalography telemetry data were collected for 236 patients with focal epilepsy. After significant data cleaning to ensure removal of epileptic activity and subsequent sleep scoring, NREM sleep data were selected for analysis. In Figure 9, the yellow star indicates the functional connectivity metrics (Figure 9A) and the values of *λ* and *α* obtained by fitting an exponential model (Figure 9B), based on in vivo data.

**Figure 9.**
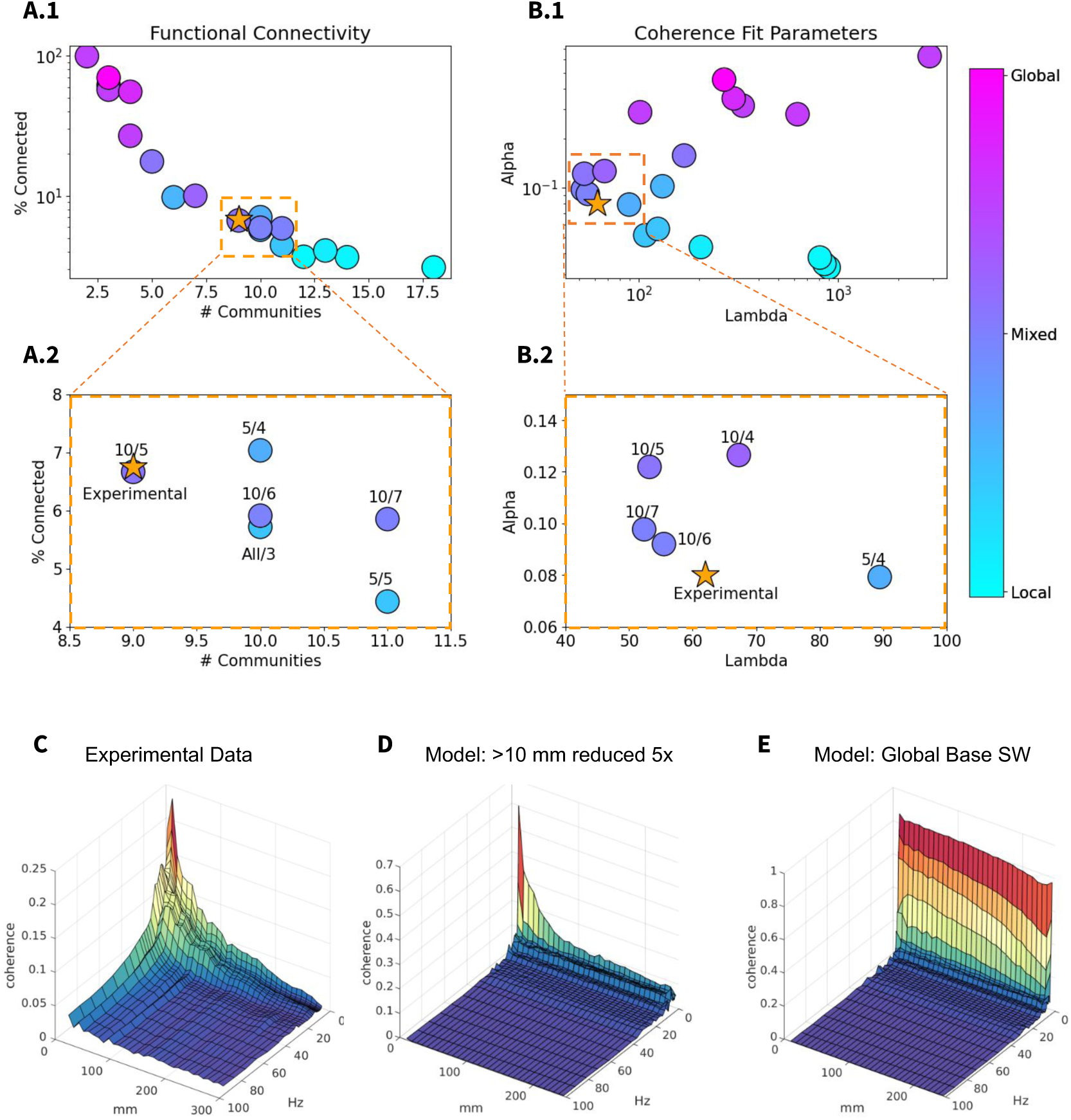
Functional Connectivity and Coherence Analysis of Experimental v.s. Model data. Coherence analysis was performed on all data sets according to the procedure described in Coherence Analysis. A) The parcel-to-parcel coherence matrices are first averaged in the slow wave frequency band and scaled by parcel-to-parcel distances to determine functional connectivity via percent of connected parcels and number of communities (determined by Louvain community detection). B) Full (unaveraged, unscaled) coherence matrices are then fitted with an exponential function to determine the full shape of the coherence landscape across distance and frequency. The resultant first (0.5 Hz) Lambda and mean Alpha in the slow wave frequency range (<2 Hz) are taken to describe the shape of the coherence landscape, and plotted in (B). (A.1 and B.1) show zoom-ins of each respective plot; the labels show the radius and factor of strength reduction - e.g., 10/5 indicates a model where connections longer than 10 mm are reduced by a factor of 5. This shows that the models with primarily mixed slow waves are the closest fit to experimental results across all 4 metrics. Full 3D coherence plots across distance and frequency are shown for the experimental data (B), the 10mm / 5 model (C), and the global slow wave base model (D), showing that the global slow wave model has a fundamentally different shape (lacking a dependence of coherence on distance) and much higher levels of coherence at low frequencies

#### Model coherence

To perform a comparable model analysis, we first averaged all the cell voltages per parcel (all layers, excitatory and inhibitory neurons) and added noise with signal-to-noise ratio (SNR) = 5 to mimic the signal that would be recorded by a depth electrode, as in experimental data (***Ball et al., 2009***; ***Suarez-Perez et al., 2018***). We then performed coherence analysis in an identical manner to the experimental data. After the full coherence matrices were computed, they were further masked to fit the sparsity of the experimental data before metrics of functional connectivity were computed.

The functional connectivity metrics reported the total percent of connected parcels, and the number of communities as determined by Louvain community detection (Figure 9A). Global models were highly connected, showing a low number of distinct communities. Conversely, local models showed very sparse connectivity, with multiple distinct communities detected. Mixed models showed moderate values on both scales. The experimental data point can be seen to fall in the middle of the distribution, surrounded by mixed models.

We then compared the exponential fits of the full coherence landscapes in the SO range (less than 2 Hz) using the mean *α* and the first *λ* value (at 0.5 Hz); these two describe the peak coherence and rate of decay. When plotted against each other (Figure 9B), all the mixed state models revealed intermediate *α* and low *λ* values. The experimental data point can be seen to fall in close proximity to the mixed state models. In particular, models in which connections longer than 10 mm were scaled down by a factor of 5, 6, or 7 were consistently very close to the experimental data point; this is also true for the model where connections longer than 5 mm are scaled down by a factor of 4 (Figure 9A.2,B.2). We found that the global slow wave model, that was initially set as a baseline model, was very far off. Note that different metrics identified slightly different variations of the models as the best match to the data (compare identified regions in Figure 8 and Figure 9); nevertheless, we found high overlap between different measurements, all indicating that the models generating mixed states are in good agreement with in vivo data.

Figure 9C,D,E shows coherence profiles obtained for in vivo data, selected mixed states model (model with a greater than 10 mm reduction, reduced 5-fold) and baseline global state model. Both the experimental data and the model displaying mixed local/global slow waves exhibit a sharp peak of coherence at low frequencies, which quickly tapers off with increased distances between parcels (Figure 9C,D). Notably, this pattern is not observed in the global slow wave baseline model (Figure 9E), where only a slight drop in coherence is seen as the distance between parcels increases. This is expected, given the unrealistic levels of slow wave synchrony across the entire network in this model. Additionally, the global slow wave model shows very high levels of coherence (nearing 1, indicating perfect coherence) at low frequencies.

In summary, these results suggest that mixed states models, particularly those with a 5-10 mm radius of stronger local connections, are the best match for experimental data.

## Discussion

Traditionally, brain states have been categorized into wakefulness, slow-wave sleep (SWS), and rapid eye movement (REM) sleep, assuming their global occurrence across brain regions. How-ever, recent large-scale neural recordings have revealed the rich heterogeneity of neural dynamics on a brain-wide scale, demonstrating that sleep and wake signatures can coexist in different brain regions, influencing behavior with spatial and temporal specificity. While local circuit mechanisms of slow waves — alternations of positive/negative EEG/LFP waves, also called slow oscillation (SO) — have been well described, the factors leading to complex heterogeneous brain states with local slow waves remain unknown. Here, we present a whole-brain thalamocortical network model based on human dMRI connectivity data capable of generating awake states and sleep states with biologically realistic cortical slow waves. The model was applied to make specific predictions about the effects of long- and short-range connectivity on the spatio-temporal dynamics of the slow waves. We identified intracortical synaptic connectivity properties that allow the coexistence of local and global slow waves and proposed mechanisms for transitions between local sleep and global uniform sleep states. The model predictions were compared to human data to validate and fine-tune the models.

A prominent example of brain states heterogeneity is local sleep (***Vyazovskiy et al., 2011***; ***Hung et al., 2013***; ***Quercia et al., 2018***; ***Engel et al., 2016***; ***van Kempen et al., 2021***). During wakefulness, local groups of cortical neurons tend to fall silent for brief periods, as they do during SWS. Further- more, during behavioral sleep, transient local wake-like activity can intrude SWS (***Nobili et al., 2011***; ***Peter-Derex et al., 2015***). It is worth noting that SWS and REM sleep can also coexist in different cortical areas (***Soltani et al., 2019***; ***Funk et al., 2016***; ***Bernardi et al., 2018***). Additionally, the proportion of time allocated to wakefulness, SWS, and REM states varies significantly across cortical areas (***Soltani et al., 2019***; ***Nazari et al., 2023***), highlighting the diverse predisposition of different regions to support these states.

From a mechanistic perspective, the basic need for local sleep can possibly be explained by homeostatic sleep pressure, that accumulates throughout the day, shaped by the duration and quality of preceding alertness. This pressure is evident in the intensity of SWS in the cortex, indicating the brain’s need for recuperation (***Wilckens et al., 2018***). Homeostatic sleep pressure can influence cortical functions in specific areas (***Siclari and Tononi, 2017***). For example, spindles and slow waves may intensify in regions engaged in learning (***Eschenko et al., 2006***; ***Pugin et al., 2015***), suggesting that sleep pressure is regulated based on recent cognitive activities. Notably, after motor skill training, there can be a localized increase in SWS within the motor cortex during rest (***Huber et al., 2004***; ***Korman et al., 2007***; ***Hanlon et al., 2009***), and a visual perception learning task can lead to a rise in the initiation of slow waves in the lateral occipital cortex (***Mascetti et al., 2013***).

Why brain areas that are more actively involved in sensory processing during the day require more SWS is a challenging question. We can speculate that it may be related to the well-characterized function of SWS in memory consolidation (***Rasch and Born, 2013***). The slow oscillation (SO) repeatedly resets the thalamocortical network during the Down phase and temporally groups sleep spindles, which are associated with long-term potentiation (LTP)-like processes and learning, during the Up phase (***Mölle et al., 2009***; ***Rosanova and Ulrich, 2005***). These observations have led to the hypothesis that SO provides a global temporal frame within the cortex and between brain regions (e.g., hippocampus, thalamus, striatum) for offline memory processing and reactivation through the strengthening of neuronal circuits (***Isomura et al., 2006***; ***Ji and Wilson, 2007***; ***Pfaff et al., 2008***; ***Rasch et al., 2007***; ***Wierzynski et al., 2009***). Local sleep may thus enable cortical networks to augment memory consolidation in specific brain regions that need it the most.

While specific mechanisms of slow-wave sleep (SWS) generation have been explored in many local circuit models (*Timofeev et al., 2000*; *Bazhenov et al., 2002*; *Compte et al., 2003*; *Hill and Tononi, 2005*; *Destexhe, 2009*; *Crunelli and Hughes, 2010*; *Levenstein et al., 2019*), only a few very recent studies (*Cakan et al., 2022*; *Goldman et al., 2023*) have proposed simulating realistic longrange cortical connectivity to test its effect on the spatio-temporal properties of SO. While these models take an important step in utilizing global brain connectivity data to study SWS, they apply a mean-field neural-mass representation of cortical regions, which limits their ability to study how the interaction of local (within a few millimeters) and long-range connectivity affects spatiotemporal SWS dynamics.

Here, we developed a large-scale thalamocortical network model based on spiking excitatory and inhibitory neurons in thalamus and cortex, organized in 6 cortical layers and thalamic core and matrix subsystems, with connectivity based on Human Connectome Project (HCP) diffusion-MRI (dMRI) tractography-based connection probabilities and HCP-derived degree of grey-matter myelination (***Rosen and Halgren, 2021***; ***Van Essen et al., 2013***). To test the contribution of specific elements of connectivity to SO properties, we then systematically modified selected elements of the network connectivity: connection density and radius, synaptic delays and strength.

While we inferred connectivity profiles from dMRI data, predicting the strength of synaptic connections is much harder problem, especially considering that the proposed model is still a significant scaling down of the biological brain. Therefore, a substantial effort of this project was to explore the strength of connections to identify the SWS regimes that match those known from the literature and characterized in our data. We found that altering the balance of excitation and inhibition had a significant impact on the synchronization properties of slow waves. While our baseline model produced unrealistically synchronized global slow-waves, reducing the strength of "longrange" excitatory connectivity resulted in models with mixed global and local slow waves, similar to results reported in vivo (***Nir et al., 2011***; ***Funk et al., 2016***). This occurred in models where connections longer than 5-10 mm were reduced four-to five-fold compared to the baseline model, to mimic the high density and strength of local connections in vivo (***Hawkins and Ahmad, 2016***; ***London and Segev, 2001***). Interestingly, this aligns with the results presented by (***Destexhe et al., 1999***), which show an approximately 7mm radius within which slow waves are highly synchronous. Furthermore, (***Riedner et al., 2007***) and (***Vyazovskiy et al., 2007***) have reported that the occurrence of multi-peak slow waves, where each peak has a different origin, increases over the course of sleep, which was proposed to result from a gradual net decrease in the strength of cortico-cortical connections (***Vyazovskiy et al., 2009***) (however, see (***Timofeev and Chauvette, 2017***)). Human recordings (***Siclari et al., 2014***; ***Bernardi et al., 2018***) reported two types of slow waves: Type I - global and more frequently occurring earlier in sleep, and Type II - local slow waves found later in sleep. A recent study suggests that global slow waves (referred to as SO) are responsible for the consolidation of new memories, while local events (referred to as delta oscillations) lead to forgetting (***Kim et al., 2019***).

To validate the model, we compared models with different synaptic connectivity profiles to human recordings during NREM sleep. This was done via a coherence analysis to determine correspondence of signals across distance and frequency and functional connectivity analysis to evaluate percentage of parcel pairs with a coherence greater than a given threshold and the number of communities. We found the experimental data to closely match the mixed state models, i.e., models showing coexistence of local and global slow-waves. Importantly, we also found that relatively modest changes of synaptic connectivity in these models lead to the models with only local or global slow-waves. Taking these results together, our model predicts that changes of the cortical synapses strength over night, may explain more frequent occurrence of global slow waves earlier in sleep. However, a more extensive analysis of synaptic connectivity in the model and analysis of in vivo data are needed to confirm this prediction.

Several studies have revealed a link between long-range intracortical connectivity and the properties of sleep slow waves. In (***Kurth et al., 2017***), it was shown that sleep slow oscillations propagate over longer distances with increasing age, and cortical SO propagation was positively correlated with intrahemispheric myelin content. Additionally, it was found that human subjects with a steeper rising slope of the slow wave exhibited higher axial diffusivity in the temporal fascicle and frontally located white matter tracts (***Piantoni et al., 2013***). These findings suggest that the profiles of sleep oscillations reflect not only the synaptic-level dynamics of the neuronal network but also the microstructural properties of its structural foundation, the white matter tracts. Factors such as normal aging (***Tomasi and Volkow, 2012***) and Alzheimer’s Disease (AD) (***Liu et al., 2014***) impact long-range connectivity. Specifically, in patients with severe Alzheimer’s Disease, functional connectivity was notably reduced between regions separated by greater distances, and this loss of long-distance connectivity was associated with a less efficient global and nodal network topology (***Liu et al., 2014***). Alongside well-documented observations of reduced slow wave density in aging (***Mander et al., 2013***) and Alzheimer’s Disease (***Katsuki et al., 2022***), these findings align with our model predictions about the critical role of balancing local and long-range connectivity for maintaining characteristic slow-wave dynamics.

The sleep SO is hypothesized to be a cortical rhythm (***Steriade et al., 1993a***; ***Sanchez-Vives and McCormick, 2000***; ***Timofeev et al., 2000***; ***Timofeev and Steriade, 1996***). In line with these results, cortical SO were maintained in the model even when thalamic input was removed. However, other studies have shown that the normal pattern of the neocortical SO requires thalamic inputs to synchronize activity across large cortical areas (***Lemieux et al., 2014***). Here, we found that removing the thalamus reduced synchrony of slow-waves in the mixed-state (but not the global) SO model, suggesting its role in long-range synchronization of cortical activity. A deeper study into thalamus involvement in SO synchrony will be a highly relevant avenue for future experimental and modeling studies. Moreover, despite not driving the SO rhythm that constitutes the object of this study, the thalamus allows this model to capture a range of human brain dynamics beyond SO. Indeed, natural extensions of this model include investigating sleep spindles, in which the thalamus is heavily involved (***Steriade et al., 1993b***, ***1990***), or the cortical impacts of thalamic stimulation, which constitute an active field of research in neuroscience (***Neudorfer et al., 2024***; ***Schüller et al., 2024***).

Although our model design achieves high computational efficiency considering the size and complexity of the network, the number of model parameters and simulation time do not easily allow a systematic fitting of the model output to electrophysiological brain signals, as it has been done with smaller computational models (***Cakan et al., 2022***). It is possible that optimization techniques such as surrogate model optimization or simulation based inference may be used on a subset of parameters in the model with the addition of biological constraints on possible values. Future studies may wish to explore this, but care will have to be taken to maintain biological plausibility. Nevertheless, the use of realistic parameters for individual neuron dynamics in wake and sleep stages (***Bazhenov et al., 2002***; ***Krishnan et al., 2016***) as well as experimentally grounded cortical connectivity (***Rosen and Halgren, 2021***) allowed us to reproduce the essential characteristics of SWS activity in human brain.

Importantly, we have now established a biologically relevant model of network dynamics that can be used for a multitude of other applications. For example, this model could be used to study possible mechanisms of sleep disorders by testing the fit of various hypothesis-driven perturbations of the model to relevant experimental data. In that context, one particularly important application of the model is using it to reveal mechanisms of sleep alterations in aging and Alzheimer’s Disease. Beyond the realm of sleep, this model could be used to study the response of a healthy brain to various types of traumatic brain injury or neurodegenerative diseases that may alter neural connectivity. Finally, this model could further provide preliminary insight on the network effects of drugs that act on specific neurotransmitters, either globally or in a targeted location.

In summary, we present a whole-hemisphere thalamocortical network model constrained by realistic local and long range connectivity based on human dMRI data and implementing effects of neuromodulators to account for sleep-wake transitions. The model displays both a biologically realistic awake state with characteristic random neuron firing as well as Type I and Type II slow wave sleep. The critical results of our study lie in revealing the separate contributions of structural connectivity (i.e., connectivity matrix) and functional connectivity (i.e., synaptic connectivity strength): while structural connectivity can alter the characteristics of individual slow waves (synchrony, spread, participation), it is primarily the proper balance of connectivity strength between local and global connections that leads to the mixed cortical states, where periods of synchronized global SO are intermittent with the periods of asynchronous local slow-waves. Importantly, only the models generating such intermittent states show spatio-temporal SO characteristics that match human recordings. Comparing the models to empirical data further supported the fundamental role of connectivity strength in generating biophysically realistic network behavior via mixed-state models. Furthermore, the model provides both the scale and the level of detail required to investigate how large-scale sleep dynamics arise from cellular and circuit level activity.

## Methods

### Model neurons

#### Map-based cortical neurons

The model has 10,242 cortical columns uniformly positioned across the surface of one hemisphere, one for each vertex in the ico5 cortical reconstruction reported in ***Rosen et al.*** (***2019***). The medial wall includes 870 of these vertices, so all analyses for this one-hemisphere model were done on the remaining 9372 columns. Each column has 6 cortical layers (L2, L3, L4, L5a, L5b and L6). For each layer in each column, one excitatory (PY) and one inhibitory (IN) neuron were simulated. To allow for large-scale network simulations we modeled cortical neurons with a model based on difference equations (MAP) (***Rulkov, 2002***; ***Rulkov et al., 2004***) which has a number of distinct numerical advantages: the common problem of selecting the proper integration scheme is avoided since the model is already written in the form needed for computer simulations. The model parameters can be adjusted to match experimental data (***Rulkov et al., 2004***; ***Bazhenov et al., 2008***; ***Rulkov and Bazhenov, 2008***). Map-based models sample membrane potential using a large discrete time step compared to the widely used conductance-based models described by ordinary differential equations, and still capture neural responses from these models. Importantly, map-based models replicate spiking activity of cortical fast-spiking, regular-spiking and intrinsically bursting neurons (***Rulkov et al., 2004***). Variations of the map model have enabled the simulation of large-scale brain networks with emergent oscillatory activity (***Rulkov and Bazhenov, 2008***), including realistic sleep spindles (***Rosen et al., 2019***) and cortical slow oscillations (***Komarov et al., 2018***). In our study, we used the map model modification proposed by ***Komarov et al.*** (***2018***), which implemented nonlinear dynamical bias, activity dependent depolarizing mechanisms, and slow hyperpolarizing mechanisms capturing the effects of leak currents, the Ca2+ dependent K+ currents and the persistent Na+ currents, which were found to be critical for simulating up/down state transitions during SO. Parameter values were initially set to the reference values in ***Komarov et al.*** (***2018***), which were shown to produce biologically realistic neural behavior during SO, and then tuned with small variations to maintain adequate neural responses in our comparatively larger-scale network. The map model equations are described below, and all parameter values specific to this study are reported in Table 2.

**Table 1.**
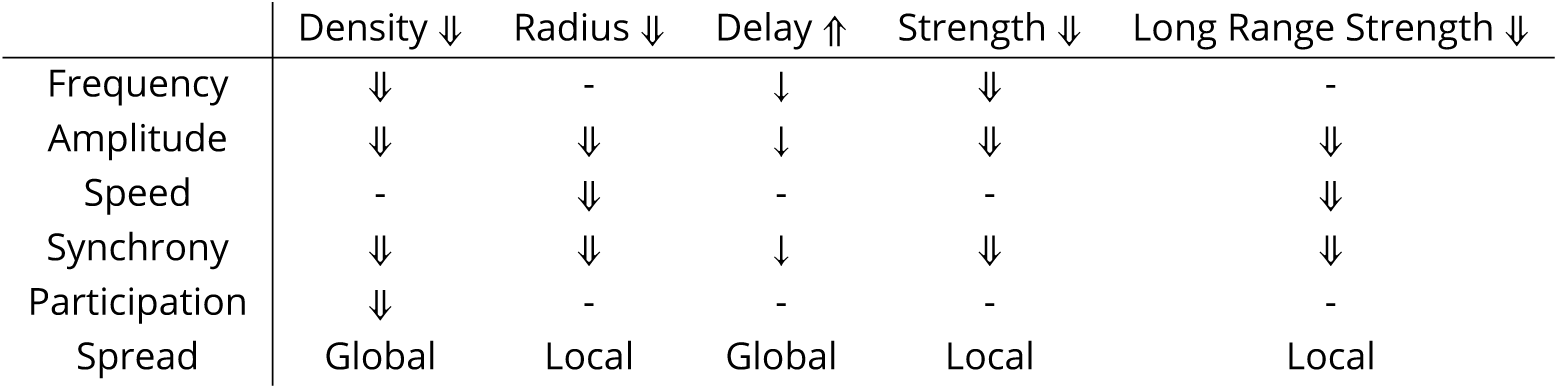
Summary of results for each network manipulation. Decreasing connection density resulted in decreased slow wave frequency, amplitude, synchrony, and spread, but slow waves remained global. Decreasing connection radius resulted in decreased slow wave amplitude, speed, participation, and spread. Increasing delays resulted in effects are seen only for manipulations far past biological plausibility, as indicated with a single down arrow for frequency, amplitude, and participation. Decreases in synaptic strength caused reduced amplitude, synchrony, frequency and participation. Finally, decreasing the strength of only long range connections resulted in traveling local slow waves with decreased amplitude, speed, and synchrony.

**Table 2.**
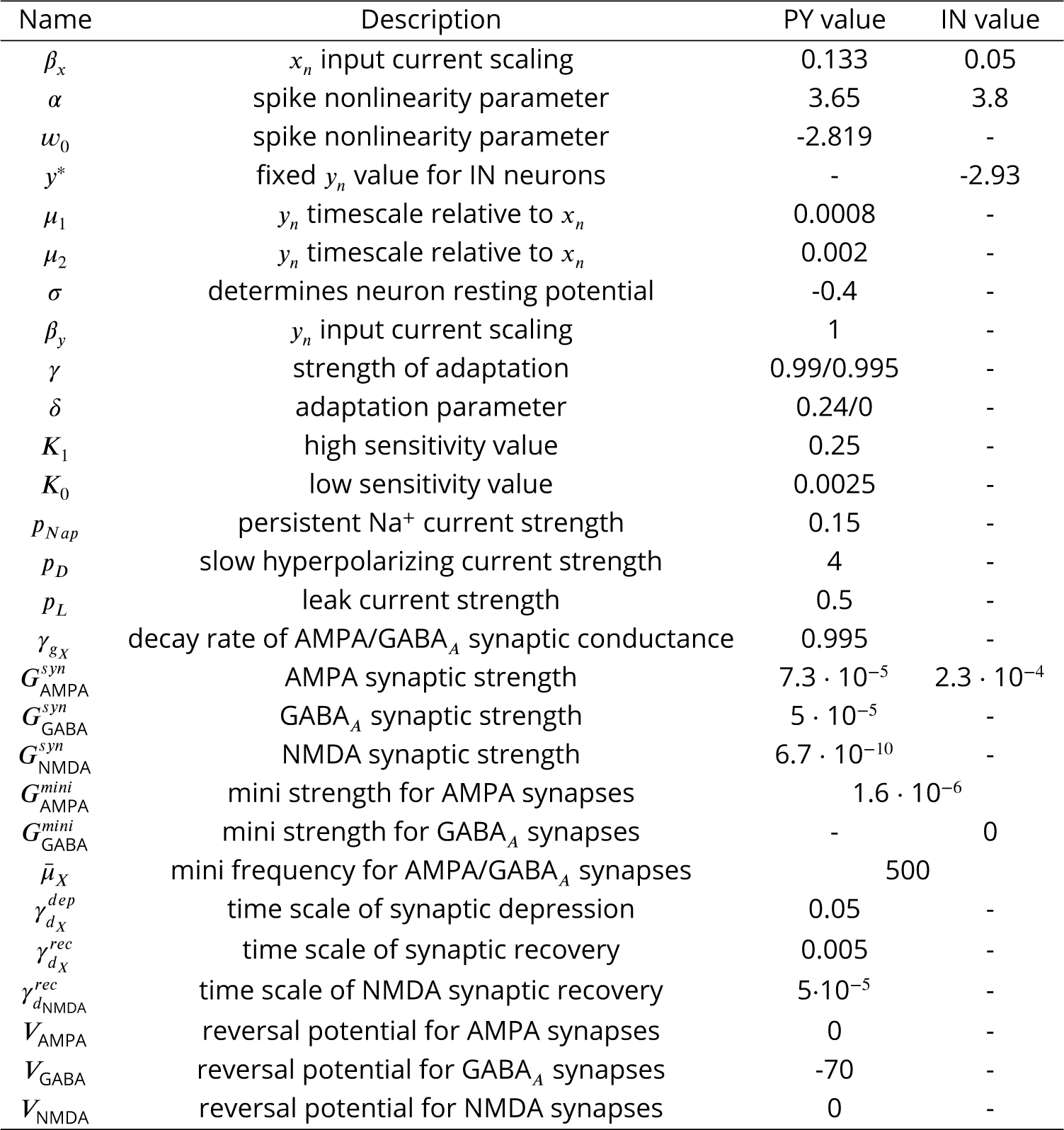
Map-based model parameter definitions and values. Parameters *γ* and *δ* may take one of the two values shown (see text). Parameters were initially set to the values in (***Komarov et al., 2018***) and then fine tuned to maintain biologically reasonable spiking behavior in the current larger-scale network. Sub-index *X* stands for "AMPA" or "GABA". For all parameters regarding synapses, the PY and IN columns denote the post-synaptic neuron

The activity of individual pyramidal PY cells was described in terms of four continuous variables sampled at discrete moments of time *n*: *x*_*n*_, dictating the trans-membrane potential *V*_*n*_ = 50*x*_*n*_ − 15; *y*_*n*_, representing slow ion channel dynamics; *u*_*n*_, describing slow hyperpolarizing currents; and *k*_*n*_, determining the neuron sensitivity to inputs. The activity of IN cells, which biologically have faster spiking dynamics, was modeled using only variable *x*_*n*_ and fixing the other variables to constant values. Dynamical PY variables evolve according to the following system of difference equations, where *n* = 1, 2, … indexes time steps of 0.5ms in size (as suggested in ***Rulkov and Bazhenov*** (***2008***)):

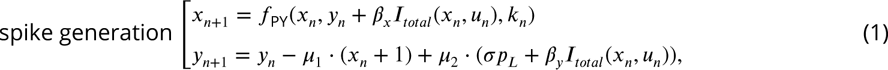

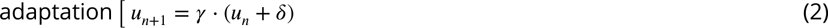

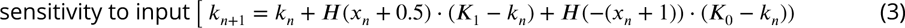

where function *H*(*w*) denotes the Heaviside step function, which takes the value 0 for *w* ≤ 0 and 1 otherwise. The reduced model for IN cells is given by

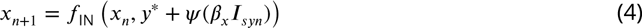

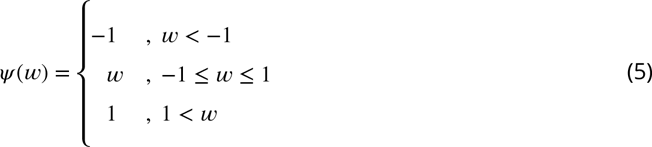

where *y*^∗^ is a fixed value of *y*_*n*_. We describe each of the system’s components below. The model’s parameters and their values are summarized in ***Table 2***.

**Spike-generating Systems** (1) and (4): Variables *x* and *y* model the fast neuron dynamics, representing the effects of the fast Na^+^ and K^+^ currents responsible for spike generation (***Hodgkin and Huxley, 1952***). The nonlinear function *f*_PY_, modified by ***Komarov et al.*** (***2018***) from ***Rulkov*** (***2002***), determines the spike waveform as

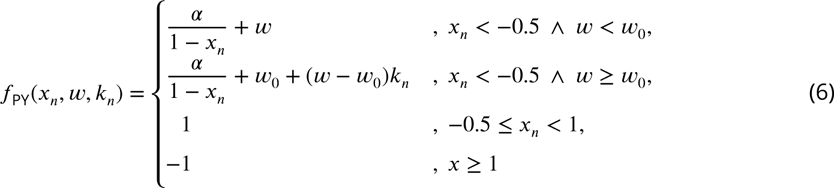

where *α* and *w*_0_ are constant parameters that control the non-linearity of the spike-generating mechanism. Parameters *μ*_1_, *μ*_2_ < 1 modulate the slower updating of variable *y* with respect to variable *x*, and parameter *σ* dictates the neuron resting potential as (1 − *σ*). The scaling constants *β*_*x*_, *β*_*y*_ modulate the input current *I*_*total*_ for variables *x* and *y*, respectively. The neuron’s membrane potential is calculated as *V*_*n*_ = 50*x*_*n*_ −15 and a spike is generated if and only if *V*_*n*_ > 0.1. In simulations, the quantity *β*_*x*_*I*_*total*_ (*x*_*n*_, *u*_*n*_) was manually prevented from reaching 0 by imposing a lower bound of 10^−4^.

For inhibitory neurons IN, function *f*_IN_ in (4) is given by

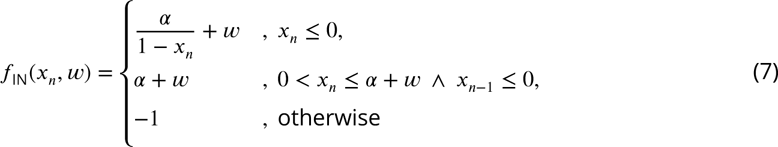

A spike is generated if and only if 0 < *x*_*n*_ ≤ *α* + *y*^∗^ + *ψ*(*β*_*x*_*I*_*syn*_) ∧ *x*_*n*−1_ ≤ 0.

**Adaptation Equation (2)**: The slow variable *u*_*n*_ represents the effects of the slow hyperpolarizing potassium currents (*I*_*D*_, explained below) reducing neural excitability over the course of the Up state and involved in Up state termination. The value of *γ* is taken as 0.99 for *x*_*n*_ < −1 and 0.995 otherwise. Parameter *δ* = 0.24 has an effect only if −0.5 < *x*_*n*_ < 1; we assume *δ* = 0 for other values of *x*_*n*_.

**Sensitivity Equation (3)**: The variable *k*_*n*_ reduces neuron sensitivity to inputs during Up states (i.e. during high spiking activity) (***Steriade et al., 2001***) by regulating the spike nonlinearity function *f*_PY_. This gives a net phenomenological representation of the combined effects of high-level synaptic activity, increase in voltage-gated hyperpolarizing currents, and/or adaptation of fast Na^+^ channels. It adopts the value *K*_0_ (low sensitivity) when voltage *x*_*n*_ crosses threshold -0.5 (i.e. Up state) and switches to *K*_1_ > *K*_0_ (high sensitivity) when *x*_*n*_ < −1 (i.e. cell leaves Up state). This prevents over-excitation and provides a physiological firing frequency (***Komarov et al., 2018***).

**Computation of currents:** The total current *I*_*total*_ is a sum of external applied DC current (*I*_DC_), synaptic inputs (*I*_*syn*_), persistent Na^+^ current (*I*_*Nap*_), slow hyperpolarizing currents (*I*_*D*_), and leak currents (*I*_*leak*_):

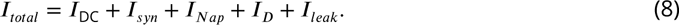

with the last three dictated primarily by *p*_*Nap*_ (maximal conductance of persistent Na^+^ current), *p*_*D*_ and *p*_*L*_ (strength of leak current), respectively:

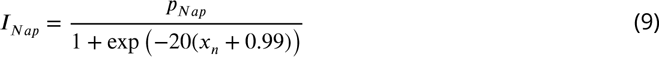

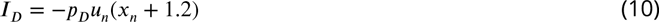

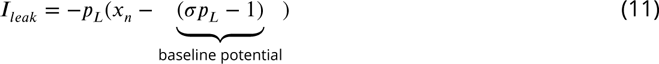

The persistent Na^+^ current contributes to initiating and maintaining Up states (***Timofeev et al., 2000***). It is modeled in *I*_*Nap*_ through a sigmoidal activation function depolarizing the neuron with strength *ρ*_*Nap*_ at high voltage values. The effect of slow hyperpolarizing currents contributing to Up state termination is represented by *I*_*D*_, which is regulated by the dynamic variable *u*_*n*_ (described above). Lastly, *ρ*_*L*_ represents the effect of potassium leak current, and here it is used to model the hyperpolarization of cortical neurons during sleep (***Bazhenov et al., 2002***). For each neuron *i*, the synaptic current 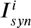 is the sum of individual AMPA, GABA_*A*_ and NMDA synaptic currents. The individual synaptic current of type *X* from neuron *j* to neuron *i* has the general form

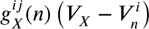

for AMPA and GABA synapses, and

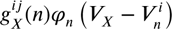

for NMDA synapses, with an additional modulator *φ*_*n*_,

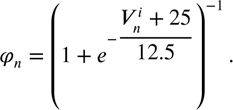

Here 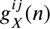 represents conductance at time *n* for directed connections from neuron *j* to neuron *i*. Dynamic synaptic conductances were described by the first-order activation schemes:

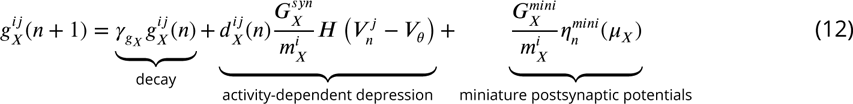

with *d*(*n*) governed by

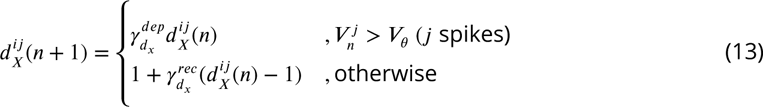

for AMPA and GABA_*A*_ and

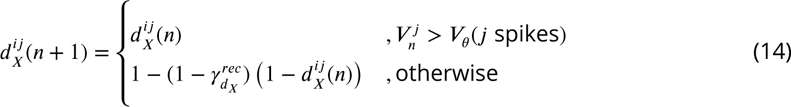

for NMDA type synapses. Here, **γ*_*gX*_* determines the decay rate of synaptic conductance and 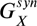 regulates the overall synaptic strength. To keep total synaptic input per cell constant, coefficients 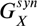 and 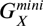 are scaled by the number 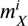 of synapses of type *X* incoming to neuron *i*. The depression variable *d*_*X*_ is dependent on whether the pre-synaptic neuron *j* spikes at time *n* or not, that is, if the voltage of neuron *j* surpasses a threshold *V*_*θ*_. Lastly, miniature post-synaptic potentials are only present in AMPA and GABA synapses. Variable 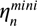 is a spontaneous event from a random Poisson process with mean frequency *μ*_*X*_, and 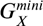 is the constant by which conductance increases in an event 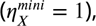 which produces miniature post-synaptic potentials ("minis") in neuron *i* (***Stevens, 1993***). Following ***Bazhenov et al.*** (***2002***), the frequency *μ*_*X*_ is modulated in time to account for the reduction in mini frequency after Up states (***Timofeev et al., 2000***), so that

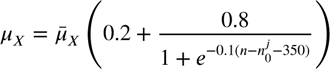

where 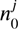 is the time of the most recent pre-synaptic spike. This modulation reduces the mini rate to 20% of its maximal value 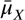 after a pre-synaptic spike, and then allows recovery to 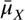 with time.

#### Hodgkin-Huxley thalamic neurons

The thalamus was modeled using a network of core (specific) and matrix (non-specific) nuclei, each consisting of thalamic relay (TC) and reticular (RE) neurons. We simulated 642 thalamic cells of each of these four types. TC and RE cells were modeled as single compartments with membrane potential *V* governed by the Hodgkin-Huxley kinetics, so that the total membrane current per unit area is given by

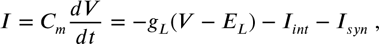

where *C*_*m*_ is the membrane capacitance,*g*_*L*_ is the non-specific (mixed Na^+^ and Cl^−^) leakage conductance, *E*_*L*_ is the reversal potential. These parameters (***Table 3***) were unchanged from (***Bazhenov et al., 1998a***), where they were determined based on previous modeling studies grounded in experimental observations (***McCormick and Huguenard, 1992***; ***Destexhe et al., 1994***). Specifically, thalamic reticular and relay neurons included intrinsic properties necessary for generating rebound responses which were found to be critical for spindle generation (***Bazhenov et al., 2000***; ***Destexhe et al., 1996***).Furthermore, *I*_*int*_ is a sum of active intrinsic currents, and *I*_*syn*_ is a sum of synaptic currents.

**Table 3.**
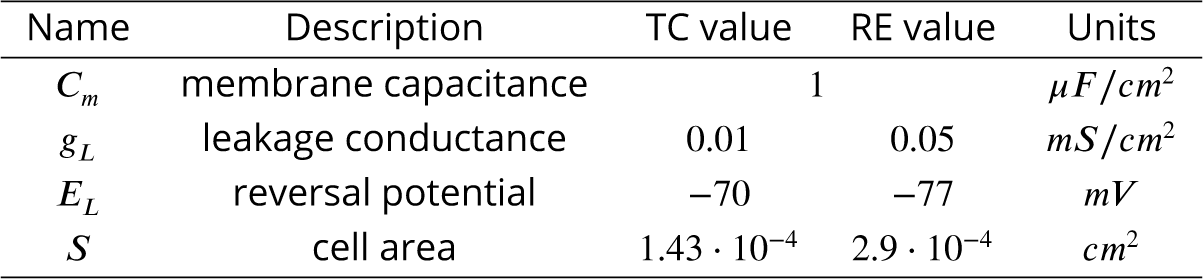
Parameter definitions and values for conductance-based models are based on previous modeling of single-cell dynamics (***Bazhenov et al., 1998a***, ***2002***; ***McCormick and Huguenard, 1992***; ***Destexhe et al., 1994***)

Intrinsic currents for both RE and TC cells included a fast sodium current, *I*_*Na*_, a fast potassium current, *I*_*K*_, a low-threshold Ca^2+^ current, *I*_*T*_, and a potassium leak current, *I*_*KL*_ = *g*_*KL*_(*V* − *E*_*KL*_). For TC cells, an additional hyperpolarization-activated cation current, *I*_*h*_, was also included. The expressions and parameter values for each of these currents are given in ***Bazhenov et al.*** (***1998a***, 2002); ***Krishnan et al.*** (***2018***); ***Rosen et al.*** (***2019***).

Synaptic currents (GABA, NMDA and AMPA) were modeled by first-order activation schemes (***Destexhe et al., 1994***), using the general form

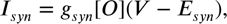

where *g*_*syn*_ is the maximal conductance, [*O*](*t*) is the time-dependent fraction of open channels, and *E*_*syn*_ is the reversal potential.Our previous studies show specific equations for all synaptic currents (***Bazhenov et al., 2002***; ***Chen et al., 2012***), as well as a detailed description of O(t) based on first-order activation schemes (***Bazhenov et al., 1998b***; ***Timofeev et al., 2000***).

#### Wake to sleep transition

The transition from wake to slow wave sleep stage was accomplished via tuning of several key parameters. The primary tunable parameters of the map model cortical neurons are *p*_*D*_ and *p*_*L*_, which control the strength of the slow hyperpolarizing currents *I*_*D*_ (similar to a calcium current) and the potassium leak currents *I*_*leak*_ as described in Equations (10) and (11). Both of these parameters significantly increased from wake to sleep. We further increased the strength of AMPA and GABA synaptic currents, as well as the strength of TC potassium leak currents *g*_*KL*_, similar to the changes described in ***Krishnan et al.*** (***2016***) (see ***Table 4***).

**Table 4.** Parameter changes between sleep and wake stages

| Parameter | Wake | Sleep |
| --- | --- | --- |
| $p_L$ | 0.15 | 0.5 |
| $p_D$ | 1.6 | 4.0 |
| $G_{AMPA}^{syn}$ (PY→PY) | $2 \cdot 10^{-5}$ | $7.3 \cdot 10^{-5}$ |
| $G_{AMPA}^{syn}$ (PY→IN) | $6.3 \cdot 10^{-5}$ | $2.3 \cdot 10^{-4}$ |
| $G_{GABA}^{syn}$ (IN→PY) | $3 \cdot 10^{-5}$ | $5 \cdot 10^{-5}$ |
| $g_{KL}$ (TC) | 0.007 | 0.012 |
| $g_{KL}$ (RE) | 0.022 | 0.015 |

### Network Structure

Our whole hemisphere computational model has realistic local and long-range cortical connectivity. Connections within the column follow those described in the canonical cortical circuit (Figure 1A). Additionally, connections within 0.1 mm are strengthened by a factor of 5 to mimic in-vivo increased strength of proximal synapses (***London and Segev, 2001***; ***Hawkins and Ahmad, 2016***). Inhibition is local: IN cells project only to PY cells in the same layer, with connections only in their own column, 1st and 2nd degree neighbors. The strength of inhibitory synapses is 2 times larger within the local column than outside. Finally, miniature postsynaptic potentials on intracortical AMPA synapses are modeled as a Poisson process (as in ***Krishnan et al.*** (***2018***)).

Thalamocortical (TC) and reticular (RE) neurons are modeled using Hodgkin-Huxley dynamics (***Hodgkin and Huxley, 1952***), with 2 subtypes of each to represent matrix and core neurons. The matrix TC neurons have a 45 mm (and 80 mm) fanout radius to cortical excitatory (and inhibitory) neurons while core TC neurons have 12 mm and 13 mm radii, respectively. Within-thalamic connections have 11 mm radii. There are 642 neurons of each of these 4 types (thalamic matrix, thalamic core, reticular matrix, reticular core).

#### Hierarchically Guided Laminar Connectivity

Long-range connectivity in the model is primarily based on the cortical parcellation into 180 areas proposed by the Human Connectome Project (HCP)-MMP1.0 atlas (***Van Essen et al., 2013***; ***Glasser et al., 2016***). Each parcel was assigned a hierarchical index inversely proportional to its estimated bulk myelination (***Glasser and Essen, 2011***; ***Burt et al., 2018***). Excitatory cortical connections (PY-PY) were then split into 6 classes according to the pre- and post-synaptic hierarchical index: within the local column, within the local parcel (or between contralateral homologs), strongly (>50th percentile) or weakly (≤50th percentile) feedforward (from lower to higher hierarchical index), and strongly or weakly (>, ≤ 50th percentile respectively) feedback (from higher to lower hierarchical index). Based on previous connectivity reports (***Markov et al., 2013***; ***Rockland, 2019***), synaptic weights for these connections were scaled by the factors shown in Figure 1 D.

#### Diffusion MRI Guided Connection Probability

The connection probability was further scaled based on previous diffusion MRI (dMRI) connectivity studies (***Rosen and Halgren, 2021***). These observations reveal an exponential decay relationship between inter-parcel dMRI connectivity and fiber distance (length constant *λ* = 23.4mm, scaling parameter *β* = 0.17). In order to obtain a probability of connection at the scale of columns, approximate intercolumnar fiber distances were estimated from geodesic distances. For parcel centroids, intra-hemispheric dMRI streamline distances and their corresponding geodesic distances were found to be related to a first approximation by a linear rational function

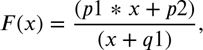

with *p*1 = 295.6, *p*2 = −2256, and *q*1 = 69.4, where *x* denotes the geodesic distance. Columnwise fiber distances were estimated by applying this function to intercolumnar geodesic distances. Thus, a distance-dependent probability of connection between columns was obtained by applying the dMRI exponential relation to column-wise fiber distances (Figure 1 E). Moreover, the residual parcel-wise dMRI connectivity (not distance related, obtained by regressing out the exponential trend) was added back to intercolumnar connections based on the columns’ parcels. This distance-independent connectivity accounts for the functional specialization of each parcel. Lastly, conduction delays were set to be proportional to fiber tract distances, with the longest connection (∼226mm) having an assigned delay of 40ms.

### Analysis

#### Modifying Network Structure

We here attempted to create the most biologically realistic model and connectivity possible. However, there is no ground truth to dictate the absolute strength of connection between two individual neurons. While rules exist in our architecture to modify relative strength based on biological connectivity, the base connection strength to be modified is a completely free (and incredibly important) parameter. We first set the base strength level to be high enough to generate Type I slow waves which quickly synchronize across the entire brain. This makes slow wave property analysis more straightforward and less prone to spatial bias when modifying further network parameters, as is the goal of this paper. Hence, we use this global Type I slow wave model for all subsequent analysis aside from analyzing the effect of connection strength itself.

In the case of modulating connection density and range, the synaptic strengths were scaled up proportionally to the loss of connections to retain sufficient activity levels in the network. The scaling mechanism works on an individual neuron basis. For each neuron, the weights of all synapses of each type (AMPA, GABA, etc.) are scaled by the total number of synapses of that type. Thus, after removing connections, the weights are divided by a smaller quantity, which results in stronger connections. This approach allows us to test the relative effects of connectivity, e.g., local vs global, without dramatic changes in the network dynamics.This allowed us to isolate the effects of patterns of connection apart from changes in total synaptic input. The ablation of connections in these trials was accomplished by setting synaptic connection strengths to 0, removing all functional connectivity.

#### Local Field Potentials, Amplitude and Frequency

In Figures 2,4,3, the local field potential (LFP) was obtained as the average voltage of Layer 2 at each timestep scaled by a factor of 20. In Figures 5,S6,7, the LFP for each local region was further smoothed using a running mean with 20ms window.

Layer LFPs were used to detect cortical Up states. A peak detection algorithm (MATLAB’s islocalmax function) was implemented for this purpose, requiring a minimum peak separation of 500ms and a minimum prominence of half the difference between the maximum and minimum values of the signal. The amplitude of each Up state was defined as the difference between the corresponding peak and the lowest point before the next peak. The inverse of each inter-peak interval was used to approximate the oscillation frequency. Average amplitudes and frequencies across all Up states of a simulation are reported in the summary plots of Figures 4,3,S7, 6.

#### Onset/offset Detection

In order to characterize how SO spread through the cortex, we identified the points in time when each neuron transitions from Down state to Up state (onset) and viceversa (offset). Our Up state detection algorithm follows ***Komarov et al.*** (***2018***), based on ***Volgushev et al.*** (***2011***): we set two thresholds, *V* ^+^ = −65mV and *V* ^−^ = −68mV (see dashed vertical lines in voltage histogram of Figure 2 B), and label any activity above *V* ^+^ as an Up state and any activity below *V* ^−^ as a Down state. This initial detection was further refined by merging any two Up states (or Down states) that were less than 100ms apart, and then removing any remaining Up state (or Down state) lasting less than 100ms, in a similar approach to ***Volgushev et al.*** (***2011***).

To identify onsets and offsets for a particular global Up state, we considered the time window between consecutive minima of the average voltage signal around that Up state (e.g. see black upward triangles in average voltage signal of Figure 2 B). For each neuron, we found the first time (if any) in this window when it reached an Up state (onset), and then the first time after the onset when it reached a Down state (offset). The minimum onset was then subtracted from all onsets so that the earliest neuron would have an assigned time of 0 and all other would have positive delays, or *latencies*. Similarly, the earliest offset was subtracted from all offsets.

In order to evaluate the synchrony of active and silent states, we constructed aggregated histograms of the onset and offset times across all Up states of each simulation. We used the standard deviation of the histograms as an measure for synchrony, with narrow histograms indicating highly synchronized global Up states.

#### Latency and participation

Latency maps are shown in Figure 2 F (also Figure S6 F, Figure 5 F and Figure 7 F) for each global Up state, representing the delay for activation of each cell after the earliest onset in that Up state. The 3D representation of latencies allows for a visual identification of the source (or sources) of each global Up state, characterized by clusters of adjacent neurons with near-zero latencies.

The percent of participating columns in each Up state was defined as the number of columns with onset delays greater than or equal to 0 over the total number of columns not belonging to the medial wall. Non-participating columns are shown in white in the latency maps for each Up state. The average percent of active columns per Up state is reported in the summary plots in Figure 3F,Figure 3F, Figure S7 and Figure 6C-D.

#### Speed of propagation

To obtain speed of propagation of each Up state, we first identified columns with onset delays under 10ms to get an approximate area of origin. We then selected the column with minimal sum of (geodesic) distances to all columns with onset delays under 10ms, excluding the medial wall. This column was identified as the origin of the Up state. We considered only an area of 150mm radius around the origin to compute propagation speed, in order to avoid interference from waves originating later in other cortical regions. A linear regression was performed between the onset delays of columns in the area and their distances to the origin. For regressions with an *R*^2^ > .25, the speed of propagation was defined as the slope of the regression line. The average propagation speed over all Up states is reported in summary plots. Speed is not reported for simulations in which there is not a linear relation between onset delay and distance from the origin, that is, simulations in which regressions for all Up states yield *R*^2^ ≤ .25.

### Experimental Data

#### Patients and intracranial recordings

Continuous stereo-electroencephalography telemetry and structural imaging were obtained for 298 patients undergoing pre-resection intracranial monitoring for phamaco-resistant focal epilepsy at University of California, San Diego Medical Center, La Jolla CA, the Cleveland Clinic, Cleveland OH, Oregon Health and Science University Hospital, Portland OR, Massachusetts General Hospital, Boston MA, Brigham and Women’s Hospital, Boston MA, the Hospital of the University of Pennsylvania, Philadelphia PA, and The University of Alabama Birmingham Hospital, Birmingham AL. These clinical data are recorded as standard of care and written informed consent, approved by each institution’s review board (The Institutional Review Board of the University of California, San Diego, The Oregon Health and Sciences University Institutional Review Board, The University of Pennsylvania Institutional Review Board, The University of Alabama, Birmingham Institutional Review Board for Human Use, The Institutional Review Board at the Cleveland Clinic Foundation, The Mass General Brigham Institutional Review Boards), was obtained from each patient before the data were used for research purposes. Recordings were collected with a Nihon Kohen Neurofax (the Cleveland Clinic), Cadwell Arc (Oregon Health and Science University Hospital), or Natus Quantum amplifier (all other clinical sites) and acquired with a sampling frequency of at least 500 Hz. After preliminary inspection, 27 patients with prior resections, highly abnormal sleep patterns, or whose data were over-contaminated with excessive epileptiform activity or technical artifacts were excluded from analysis. A further 35 patients were excluded from further analysis after channel selection and quality control procedures, described below, resulted in them having with fewer than two clean bipolar channel pairs. After quality control, 236 patients (126 female, 101 male, 9 unknown, 36.5 ± 13.3 (mean ± stdev.) years old, 22 unknown age) were included in the final analysis.

#### Channel localization and inter-channel distance

As part of standard of care, a pre-operative T1-weighted high-resolution structural MRI and interoperative CT scan were obtained for each patient. SEEG contacts were localized as previouslt described (***Dickey et al., 2022***). Briefly, the post-implant CT volume was co-registered to the pre-implant MR volume in standardized 1mm isotropic FreeSurfer space with the general registration module (***Johnson et al., 2007***) in 3D Slicer (***Fedorov et al., 2012***) and each contact’s position was determined by manually marking the contact centroid as visualized in the co-registered CT volume. Cortical surfaces were reconstructed from the MR volume with the FreeSurfer recon-all pipeline (***Fischl, 2012***). Each transcortical bipolar channel was assigned to the white-gray surface vertex nearest to the midpoint of the adjacent contacts. Spherical folding-based surface registration (***Fischl et al., 1999***) was used to map the individual subjects’ bipolar channel vertices to the standardized fsaverage ico7 (***Fischl, 2012***) and 32k_FS_LR grayordinate (***Marcus et al., 2013***) atlases as well as to parcels of the HCP-MMP parcellation (***Glasser et al., 2016***). The HCP-MMP parcel assignments of each channel were used to determine the approximate white matter fiber length of the connection between them, using the mean values for a cohort of 1,065 healthy subjects we have previously reported (***Rosen and Halgren, 2021***).

#### Telemetry preprocessing and channel selection

For each patient, an average of 185.3 ± 112.0 hours of raw telemetry were analyzed. Data were anti-aliased filtered at 250 Hz and resampled to 500 Hz. To remove line noise, a notch filter was applied at 60, 120, 180, and 240 Hz. On each electrode shaft, data were re-referenced to a bipolar scheme by taking the difference of potentials in adjacent contacts, yielding 30,769 total channels. To ensure that bipolar channels were independent, contact pairs were selected such that no two pairs shared a contact, eliminating just under half of all channels from further consideration. Peaks in the delta band (0.5-2 Hz) signal during high-delta periods were identified and the peri-delta-peak high-gamma band (70-190 Hz) envelope obtained. Transcortical contact pairs were identified by (1) having a midpoint close to the gray-matter–white-matter interface (<5 mm), (2) having a high anticorrelation between the delta-band amplitude and the high gamma band (70-190 Hz) envelope during delta-band peaks, indicative of non-pathological Down states with a surface-negative LFP deflection that quieted spiking, and (3) having high delta-band amplitude with and average delta peak of at least 40 *μ*V, indicative of robust inversion in adjacent contacts. Only contacts with 5 mm pitch were analyzed to standardize absolute bipolar amplitudes and contact pairs located deep in the white matter or in subcortical gray matter were excluded from analysis. All channels were visually inspected to ensure they contained broadly normal non-pathological waveforms such as pathological delta activity. This procedure yielded 4,903 candidate channels which underwent NREM scoring, described below, yielding 9.2 ± 7.8 hours of putatively clean scored NREM sleep. After scoring, which excluded continuous periods of artifactual or pathological signals, remaining epileptogenic activity was detected with spike template convolution and gross artifacts, with amplitudes > 300 *μ*V. Channels which contained persistent automatically detected interictal spikes or gross artifacts were removed from further analysis, yielding 3,665 channels. Periods containing interictal spikes or gross artifacts in any channel were excluded from analysis in all channels. Finally, after cross-spectral matrices were obtained for each subject as described below, channels with autospectral power more than 3 standard deviations above the pooled channelwise mean in any examined frequency were excluded, leaving 3,588 channels and 37,123 within-patient bivariate channel pairs in the final analysis.

#### NREM selection

Clean periods of sleep were first manually classified based on ultradian delta power. Within manually identified nightly periods of sleep, NREM sleep was automatically identified by the presences of slow oscillations (SOs) in a manner derived from standard clinical procedures (***Iber et al., 2007***). The density of SOs was ascertained in each 30-second frame and NREM marked for the frame if at least 35% of the frame contained SOs in at least one bipolar channel. The reduction in the number of channels and increase in the percent of frame threshold relative to the clinical standard of 20% of the frame across scalp channels was to accommodate the increased spatial heterogeneity of sleep graphoelements observed intracranially, relative to the scalp (***Mak-McCully et al., 2017***). For this purpose, slow waves were automatically detected with the method described in (***Mak-McCully et al., 2017***), briefly, consecutive delta-band zero-crossings occurring within 0.23-3 s were detected and the top 20% of peaks were retained as SOs. Only periods without detected interictal spikes or gross artifacts in any bipolar channel were included in subsequent analysis.

### Coherence Analysis

#### Mock Experimental Data

Coherence analysis was performed on both experimental and model data in an identical manner following preprocessing. The model data was preprocessed by averaging the voltages from all cells belonging to each of the 180 parcels (all 6 layers, both excitatory and inhibitory) to mimic what would be recorded by an SEEG electrode. Additionally, white gaussian noise was added to each of the 180 parcel averaged signals. This was done to mimic experimental noise picked up from both other non-neural bodily signals (i.e., muscle movements) and noise from the recording electrode itself. ***Ball et al.*** (***2009***) demonstrated that ECOG arrays show an SNR ranging from 2.22 to 4.37 in relation to eye blink movement artifacts. ***Suarez-Perez et al.*** (***2018***) further calculated the SNR of slow wave sleep Up states (signal) v.s. Down states (noise); they found low frequencies (less than 30 Hz) to have a steady SNR of 4-9, depending on the type of electrode used. At higher frequencies, the SNR dropped severely. With these studies in mind, we implemented an SNR of 5 to account for both body and electrode noise that our model data lacks.

#### Signal processing and epoch sampling

A frequency resolved map of each patient’s (or simulation’s) neural covariability was achieved by estimating the complex cross-spectral matrix among bipolar SEEG channels (for empirical data) or model parcel averages. Narrow-band signals were extracted with a series of second-order resonator filters (matlab’s iirpeak) with peaks and 3 dB attenuation bandwidths shown in Table S1. For each frequency the complex analytic signal was extracted via the Hilbert transform. Calculating the variance-covariance matrix of analytic signals yields the cross-spectral matrix. Cross-spectral matrices were normalized into magnitude-squared coherence by squaring the complex modulus of each cross-pairwise cross-spectrum and dividing by the product of the constituent autospectra.

For experimental data, an adaptive procedure was used to obtain stable representative estimates of spontaneous covariability for each patient. Continuous minutes of telemetry were iteratively sampled at random, without replacement, from artifact-free NREM sleep. Sampled minutes of telemetry were appended and prepended with 1.5 seconds of data for filtering and analytic signal extraction. These data were removed prior to covariance estimation to avoid edge artifacts. For each frequency, the coherence matrices were obtained as described above for the incoming minute, _*n*+1_Σ_*n*+1_, the cumulatively sampled telemetry, _1_Σ_*n*+1_, and, for the second sampled minute onward, the prior cumulatively sampled telemetry, _1_Σ_*n*_. The correlation matrix distance (CMD) (***Herdin et al., 2005***), was calculated between _1_Σ_*n*_ and _1_Σ_*n*+1_, and between _1_Σ_*n*_ and _*n*+1_Σ_*n*+1_. Starting with the third sampled minute, incoming minutes were rejected if the CMD between _1_Σ_*n*_ and _*n*+1_Σ_*n*+1_, was more than 3 standard deviations greater than average CMD between _1_Σ_*n*_ and _*n*+1_Σ_*n*+1_ for all previously sampled minutes. The coherence matrix was considered to have converged when the CMD between _1_Σ_*n*_ and _1_Σ_*n*+1_ was less than 1 × 10^−5^ and the cumulative cross-spectral matrix was then retained. To dilute the influence of the first sampled minute, this procedure was repeated ten times, and the mean of the resulting cross-spectral matrices was used for subsequent analyses.

#### Statistical analyses and model fitting

The effect of inter-parcel fiber distance on coherence was assessed by fitting a linear mixed-effects model: log(coherence) distance + 1 + (1|patient), which accounted for the fixed effects of the predictor variable distance and an intercept term, as well the random effect intercept term grouped by individual (this term was excluded for simulated data). Because of the natural log transformation of the response variable coherence, the linear model is equivalent to an exponential model for the untransformed distance predictor. Therefore, we report the modeled slopes and intercepts as length constants (*λ*) and scaling terms (*α*, maximum coherence), respectively. Separate fits were made for each narrowband frequency. Because the parameters of the exponential trend appeared to change at distances greater that 100 mm in experimental, especially for higher frequencies, separate models were fit to electrode pairs 0-100 mm distant, >100 mm, as well as to all pairs. Despite having twice as many parameters, the two-domain model was favored over the one-domain model by both Bayesian and Akaike information criteria for all frequencies and behavioral states. Only the 0-100 mm model parameters are reported here for both experimental and simulation data. All mixed effects linear models were fitted with matlab’s fitlme. To avoid the inherent skewness coherence coefficient distributions, coherence values were transformed into z-statistics (***Fisher, 1921***) before model fitting or averaging for display then reverted into coherence coefficients after model fitting or averaging.

Finally, we take the 0.5 Hz lambda and the mean <2 Hz alpha values to describe the key rate of decrease and max coherence in the slow wave range as the primary characteristics of each coherence landscape to be compared across experimental and model data (Figure 9).

## Acknowledgments

This research was supported by the National Science Foundation Graduate Research Fellowship Program under Grant No. DGE-2038238, and the National Institutes of Health (NIH) (grants 1R01MH117155, 1R01NS109553, 1R01MH125557). Data were provided [in part] by the Human Connectome Project, WU-Minn Consortium (Principal Investigators: David Van Essen and Kamil Ugurbil; 1U54MH091657) funded by the 16 NIH Institutes and Centers that support the NIH Blueprint for Neuroscience Research; and by the McDonnell Center for Systems Neuroscience at Washington University.

## Supplementary Information

**Figure S1.** Neuronal activity in all 6 layers during the Global SO model from Figure 2. Voltage traces in mV for two different cells are shown on the left panels, with individual spikes marked under each trace in the same color.

**Figure S2.** Activity of pyramidal, inhibitory and thalamic RE and TC neurons in baseline model in Figure 2B.Voltage traces in mV for two different cells are shown on the left panels, with individual spikes marked under each trace in the same color.

**Figure S3.** Activity of cortical cells in all layers in mixed model.Voltage traces in mV for two different cells are shown on the left panels, with individual spikes marked under each trace in the same color.

**Figure S4.** Activity of pyramidal, inhibitory, and thalamic cells in the mixed model. Voltage traces in mV for two different cells are shown on the left panels, with individual spikes marked under each trace in the same color.

**Figure S5.** Activity of pyramidal and inhibitory neurons in the Activity by Layer and Cell Type: Mixed local/global model in Figure S4, with the thalamus isolated from the cortex. Voltage traces in mV for two different cells are shown on the left panels, with individual spikes marked under each trace in the same color.

**Figure S6.** Local activity for *P* = 0.1, with 90% connections removed. A) Ten cortical areas with a 5mm radius, that were used to calculate local dynamics. B) Average membrane voltage of layer II neurons, as in Figure 3e.2. C-E) For each region in (A), subpanels show: (C) the single-cell voltage for two neurons in the area, (D) the local field potential (LFP) for the 5mm area, and (E) heatmap of individual voltages of all neurons in the area. Up states are largely synchronized across all 5 regions. F) Latency map for each Up state in the *P* = 0.1 simulation. Even with very sparse connectivity, Up states spread to the whole cortex. Participation was reduced uniformly to about 70% compare to nearly 100% participation when all connections are present (compare to Figure 2D)

**Figure S7.** Effect of Synaptic Delay, either via setting a uniform delay (A) or scaling the max delay (B) on SO dynamics. (A and B) Summary plots of frequency, amplitude, and onset / offset spread. Decreasing the max scaled delay to 0.1 ms (from 40 ms) has no effect, while increasing the max scaled delay up to 1 second shows only minor changes in frequency frequency and amplitude but notably increased onset/offset synchrony. Low uniform delays similarly showed no effect, with loss of amplitude and synchrony only seen at 30 ms (most delays were previously under 2 ms).

**Figure S8.** Activity maps and graph properties for the full-connectivity, Global SO network in Figure 2. Rows from top to bottom show (1) the percent of simulation time that each pyramidal cell in layer 2 spends in an Up state; (2) the mean onset/offset delay across Up states for each pyramidal cell in layer 2; (3) the number of strongly/weakly connected components in the structural connectivity graph, constructed with each cortical column as a node and each directed edge weight as the number of synaptic connections from one column to another; (4) the normalized in/out-degree for each node in the graph; and (5) the normalized in/out-degree centrality for each node in the graph, defined as the number of synaptic connections to/from that column (see Graph Properties for details). Initiation zones (low mean onset delay) at the front and back and have generally lower in-degree and out-degree than areas in the middle.

**Figure S9.** Activity maps and graph properties for the *R* = 2.5mm network in Figure 5. Rows from top to bottom show (1) the percent of simulation time that each pyramidal cell in layer 2 spends in an Up state; (2) the mean onset/offset delay across Up states for each pyramidal cell in layer 2; (3) the number of strongly/weakly connected components in the structural connectivity graph, constructed with each cortical column as a node and each directed edge weight as the number of synaptic connections from one column to another; (4) the normalized in/out-degree for each node in the graph; and (5) the normalized in/out-degree centrality for each node in the graph, defined as the number of synaptic connections to/from that column (see Graph Properties for details). Regions with high percent time active have high in/out-degree and centrality

**Table S1.** Peaks and 3 dB attenuation bandwidths for the second-order resonator filters used to extract narrow-band signals.

**Video S1.** The base global slow wave sleep model over 30 seconds of simulated time. The top plot shows Layer 2 cell voltages across the cortex, while the bottom plot shows the average voltage trace.

**Video S2.** The slow wave sleep model with a connection density of P = 0.1 over 30 seconds of simulated time. The top plot shows Layer 2 cell voltages across the cortex, while the bottom plot shows the average voltage trace.

**Video S3.** The slow wave sleep model with a connection range of R = 2.5 mm over 20 seconds of simulated time. The top plot shows Layer 2 cell voltages across the cortex, while the bottom plot shows the average voltage trace.

**Video S4.** The slow wave sleep model with connections longer than 10mm reduced 5-fold over 105 seconds of simulated time. The top plot shows Layer 2 cell voltages across the cortex, while the bottom plot shows the average voltage trace.

