## Supplemental Information for "Emergent effects of synaptic connectivity on the dynamics of global and local slow waves in a large-scale thalamocortical network model of the human brain"

### Supplementary Information

#### Activity by Layer and Cell Type: Global SO model

Figure S1 shows the activity of all 6 modeled layers baseline SO simulation in Figure 2. Activity is largely synchronized across layers. Layer 4 is consistently less active than other layers, which is likely a result of L4 PY cells having within-column connections from exclusively one other layer (L6, see Figure 1E)

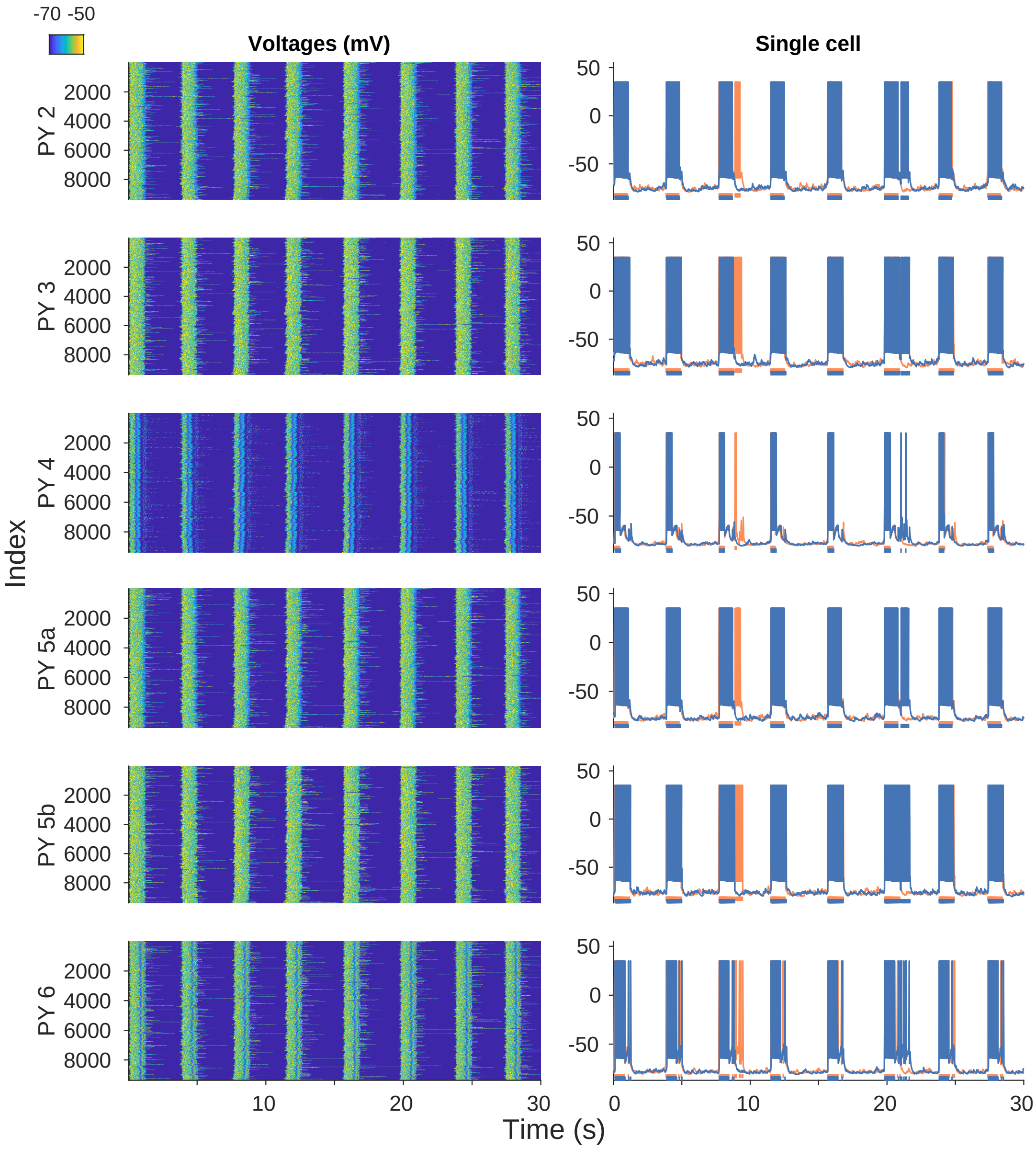


**Figure S1.** Neuronal activity in all 6 layers during the Global SO model from Figure 2. Voltage traces in mV for two different cells are shown on the left panels, with individual spikes marked under each trace in the same color.

Figure S2 shows the activity of modeled core and matrix thalamocortical (TC, TCa) and reticular (RE, REa) cells as well as inhibitory (IN) cells.

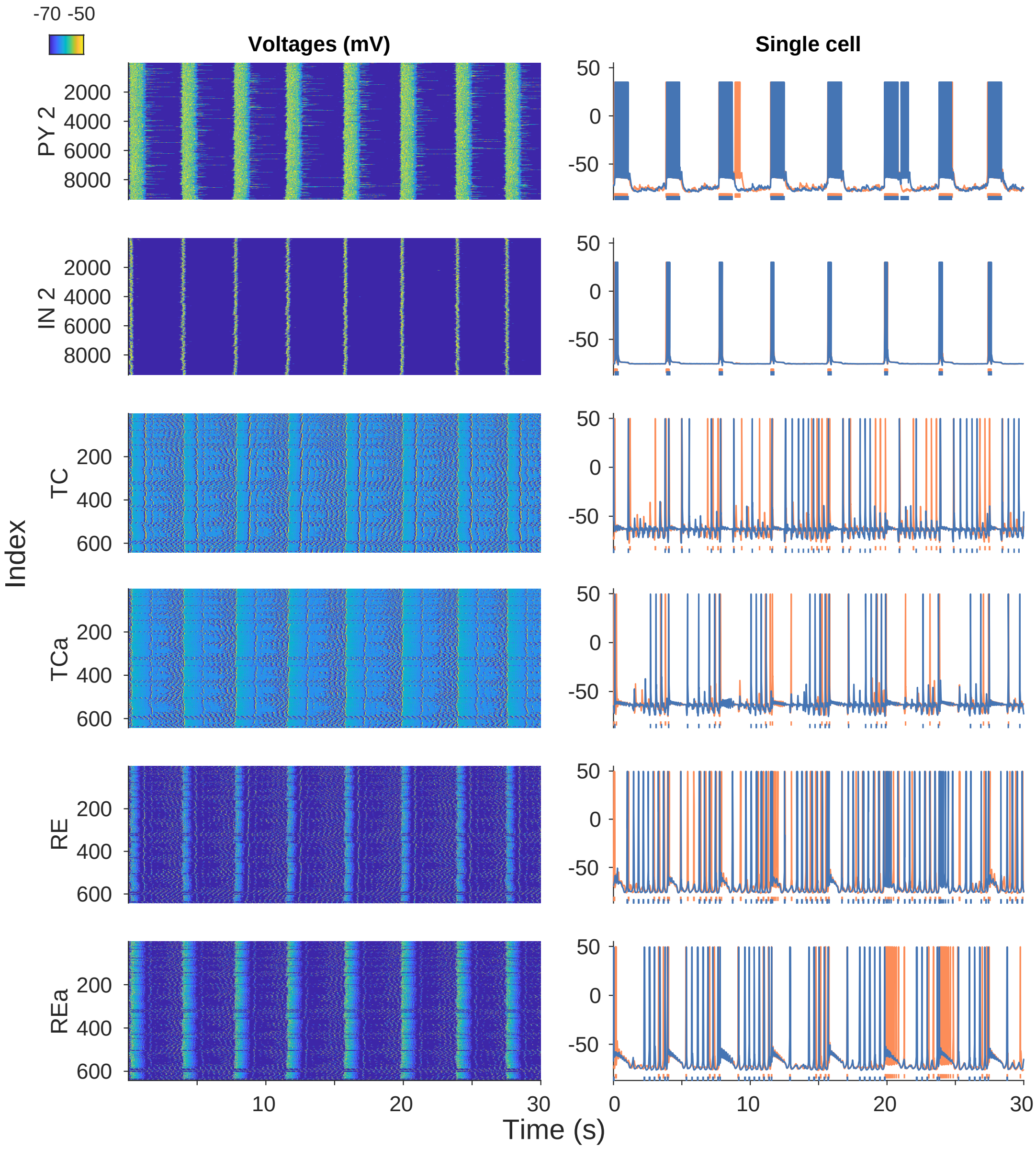


**Figure S2.** Activity of pyramidal, inhibitory and thalamic RE and TC neurons in baseline model in Figure 2B.Voltage traces in mV for two different cells are shown on the left panels, with individual spikes marked under each trace in the same color.

### Activity by Layer and Cell Type: Mixed local/global model

Figure S3 shows all layers of cortical cells in the mixed model (Local vs Global Slow-Waves) with global and local Up states. Similar to the Activity by Layer and Cell Type: Global SO model, layers are largely synchronized, with a relatively less active L4.

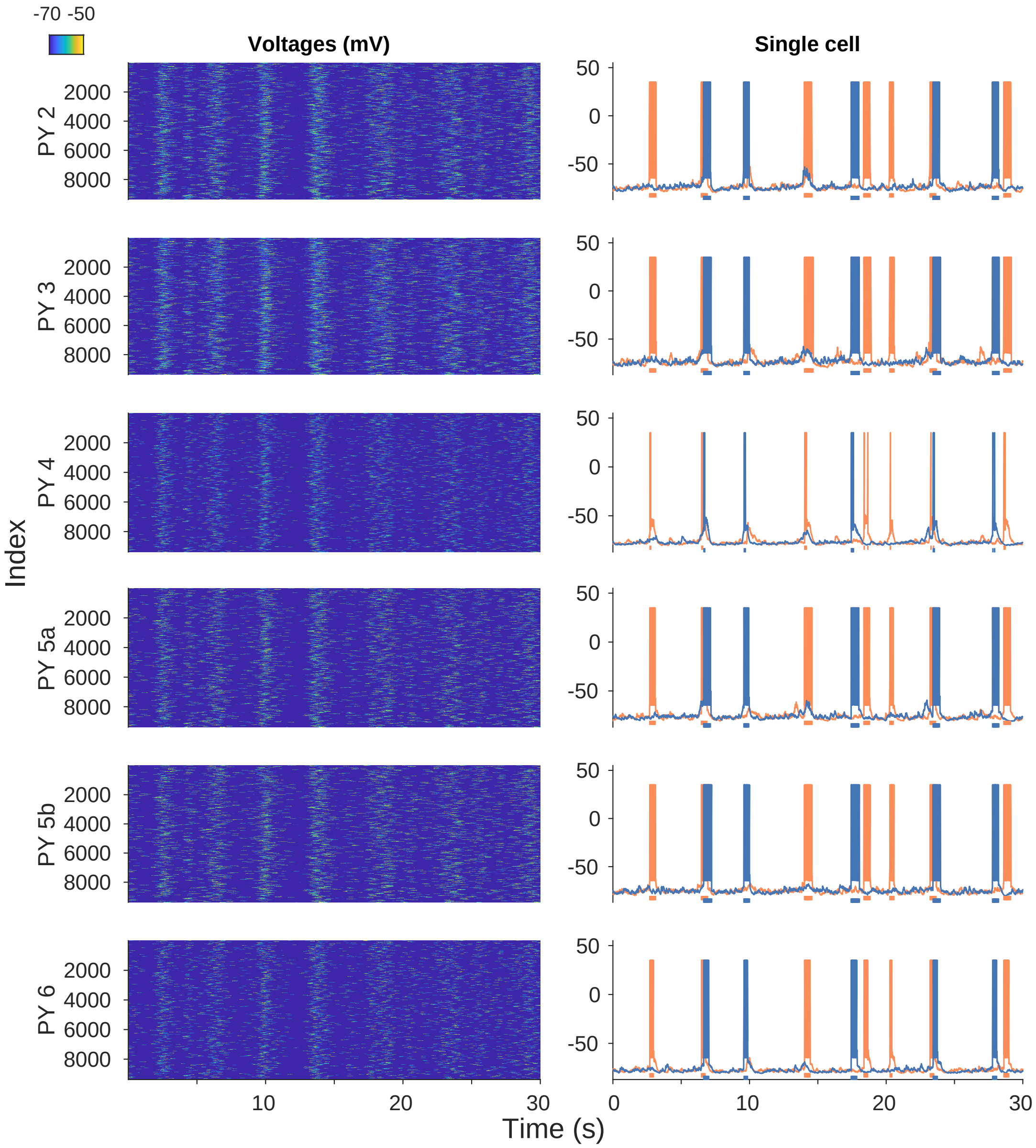


**Figure S3.** Activity of cortical cells in all layers in mixed model.Voltage traces in mV for two different cells are shown on the left panels, with individual spikes marked under each trace in the same color.

1364 Figure S4 additionally shows the activity of inhibitory and thalamic core and matrix cells (TC, 1365 TCa, RE and REa).

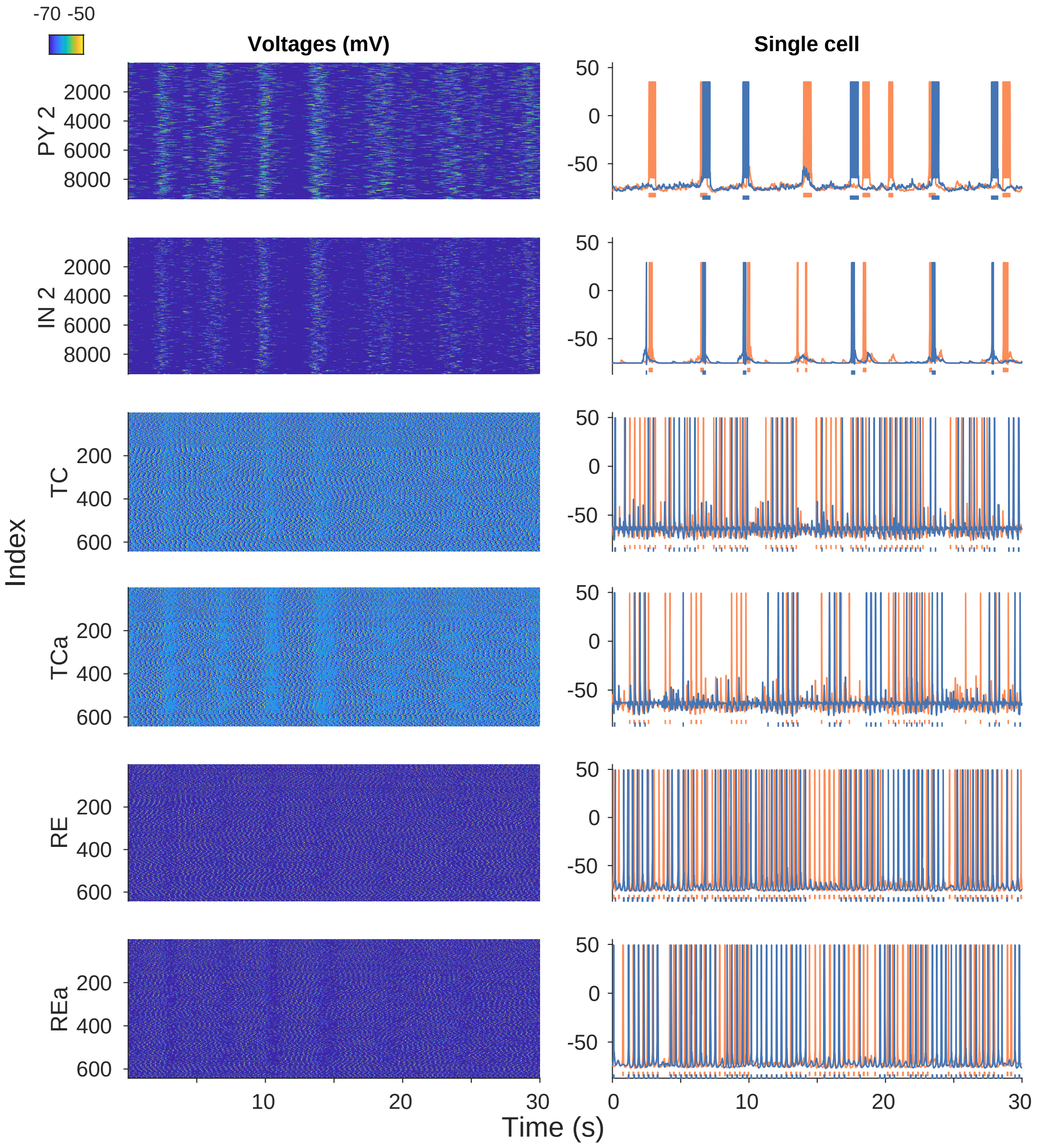


**Figure S4.** Activity of pyramidal, inhibitory, and thalamic cells in the mixed model. Voltage traces in mV for two different cells are shown on the left panels, with individual spikes marked under each trace in the same color.

### Effect of thalamus on SO synchronization

Figure S5 shows activity of pyramidal cell and inhibitory INs in the Activity by Layer and Cell Type: Mixed local/global model when the thalamus and cortex are isolated. SO activity is maintained in cortical cells, but less synchronized than in simulations including the thalamus (compare to Figure S4). Thalamic cells (not shown) remain inactive in the absence of cortical inputs. On the other hand, removing the thalamus in the Activity by Layer and Cell Type: Global SO model does not produce an appreciable effect, likely because the impact of thalamic inputs diminishes relative to the elevated cortical synaptic weights, which are high enough to produce synchronous Up states.

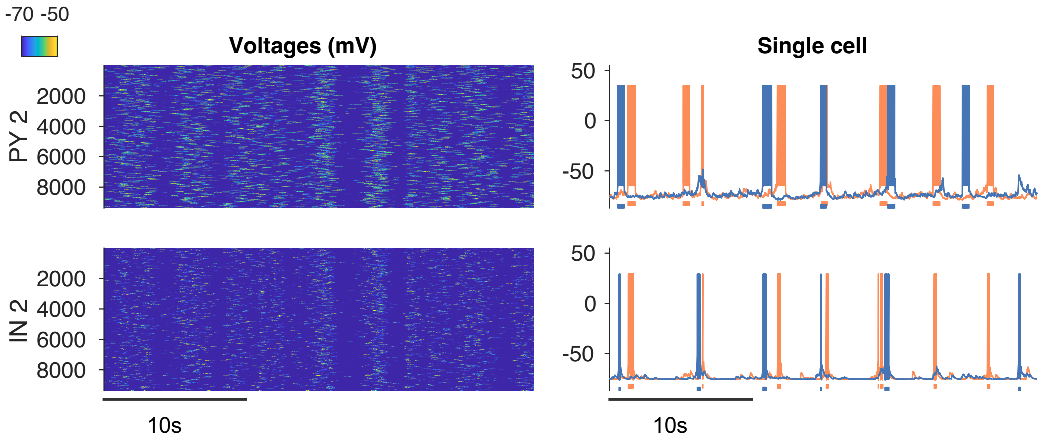


**Figure S5.** Activity of pyramidal and inhibitory neurons in the Activity by Layer and Cell Type: Mixed local/global model in Figure S4, with the thalamus isolated from the cortex. Voltage traces in mV for two different cells are shown on the left panels, with individual spikes marked under each trace in the same color.

### Effects of connections density, range and delays

Figure S6 shows further details into the slow-wave dynamics in the case of extreme density loss ($P = 0.1$), zooming into ten cortical regions with a 5mm radius each (Figure S6A). Single cells and LFPs in different regions throughout the cortex (Figure S6C-D) showed synchronized Up states, although many neurons were not active during individual Up states (Figure S6C, E). The local heterogeneity in activation patterns induced by the extreme sparsity of connections is evident in Figure S6E, which shows many silent neurons in the immediate vicinity of active cells. This generated the extremely granular participation seen in the latency maps (Figure S6F). Even though each Up state encompassed cells in the whole cortex (high spread), a large fraction of cells distributed through the entire cortex did not participate in slow-waves (decreased participation). This model can further be seen in Supplemental Video S2. Initiation was diffused with respect to the baseline simulation, to the point where a single meaningful estimate of propagation speed for each Up state could no longer be computed.

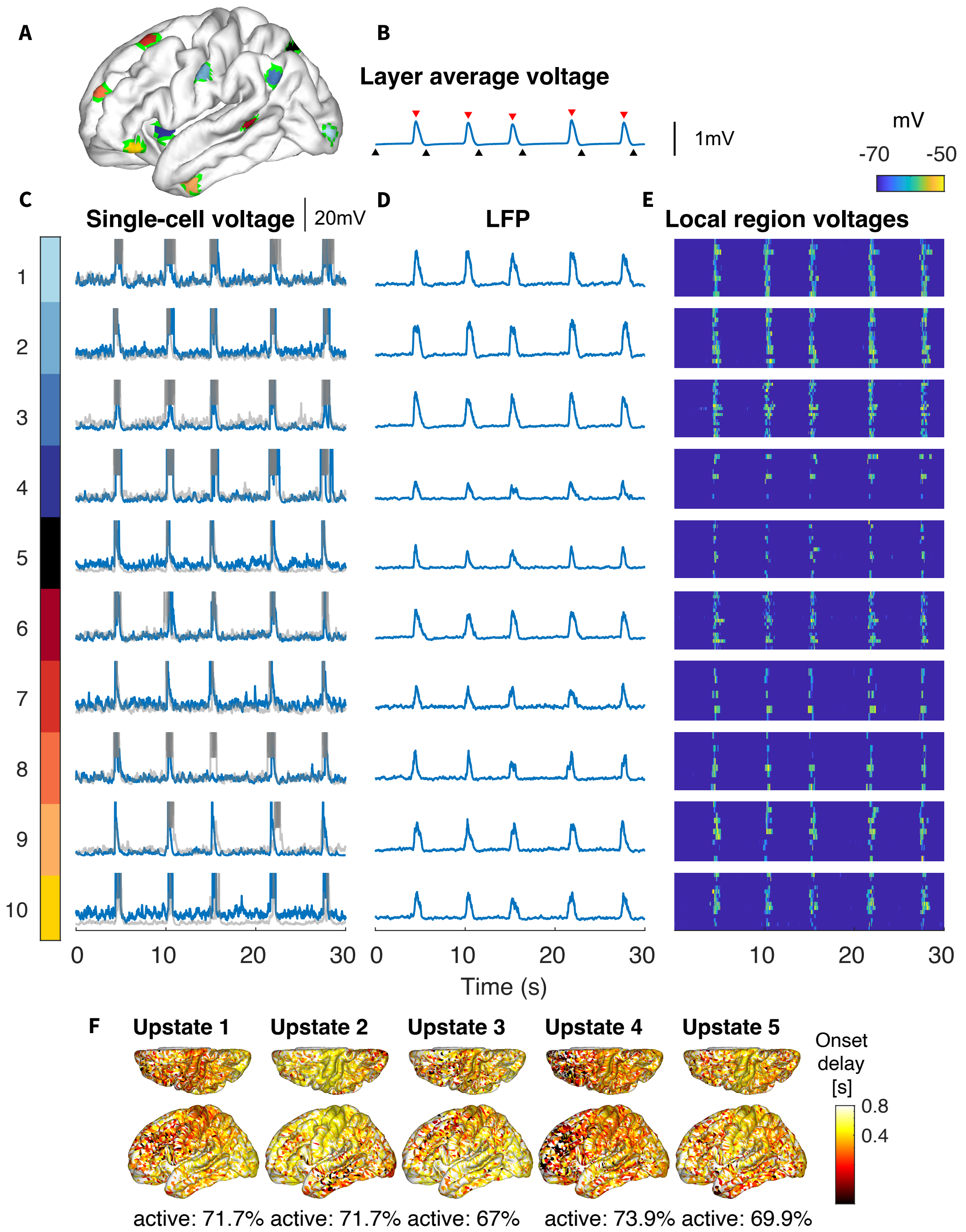


**Figure S6.** Local activity for $P = 0.1$, with 90% connections removed. A) Ten cortical areas with a 5mm radius, that were used to calculate local dynamics. B) Average membrane voltage of layer II neurons, as in Figure 3e.2. C-E) For each region in (A), subpanels show: (C) the single-cell voltage for two neurons in the area , (D) the local field potential (LFP) for the 5mm area, and (E) heatmap of individual voltages of all neurons in the area. Up states are largely synchronized across all 5 regions. F) Latency map for each Up state in the $P = 0.1$ simulation. Even with very sparse connectivity, Up states spread to the whole cortex. Participation was reduced uniformly to about 70% compare to nearly 100% participation when all connections are present (compare to Figure 2D)

Figure S7 shows the effect of modifying synaptic delays on network behavior. Delays were changed in two ways: (1) by setting a maximum delay and scaling all other delays with distance and (2) by imposing a fixed delay value for all connections and varying this value (see Connection Delay for details)

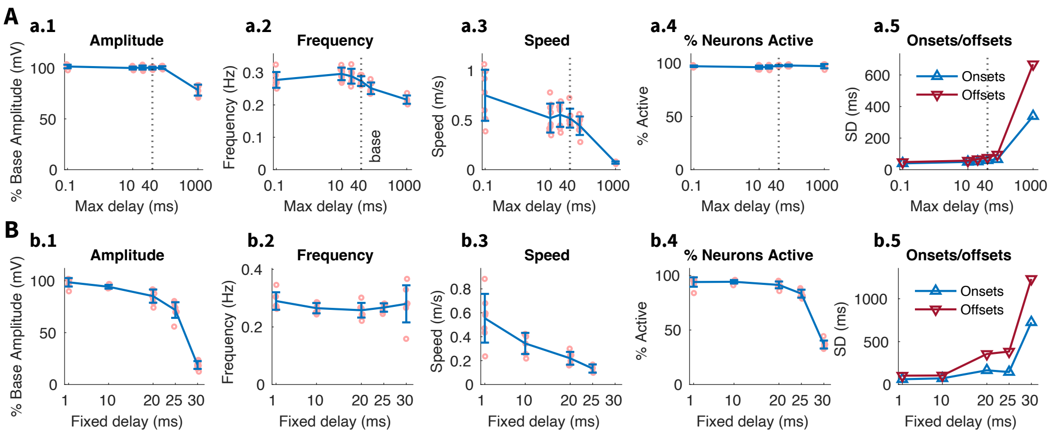


**Figure S7.** Effect of Synaptic Delay, either via setting a uniform delay (A) or scaling the max delay (B) on SO dynamics. (A and B) Summary plots of frequency, amplitude, and onset / offset spread. Decreasing the max scaled delay to 0.1 ms (from 40 ms) has no effect, while increasing the max scaled delay up to 1 second shows only minor changes in frequency frequency and amplitude but notably increased onset/offset synchrony. Low uniform delays similarly showed no effect, with loss of amplitude and synchrony only seen at 30 ms (most delays were previously under 2 ms).

### Graph Properties

Figure S8 and Figure S9 show activity maps and graph properties of the networks for the Global slow wave simulation in Figure 2 and for the restricted range $R = 2.5mm$ simulation in Figure 5. Activity characteristics (percent time active, mean onset/offset delays) are shown for pyramidal neurons in cortical layer 2 only for 30 seconds of simulation time, as in Slow Wave Characterization and Connection Range. The percent time active for each neuron is defined as the fraction of the simulation time that the neuron spends in an Up state (above $-65$mV, see Latency and participation). The mean onset/offset delay for each cell is an average of its onset/offset values across Up states (see Latency and participation for details on onsets/offsets).

The graph properties (strongly/weakly connected components, in/out degree, and centrality) are based on structural connectivity only, independent of activity during simulations. A weighted directed graph is constructed for each network, where each column is a node and the edge from node $i$ to node $j$ has an associated weight equal to the number of synaptic connections from column $i$ to column $j$. For each graph, Figure S8 and Figure S9 show: the number of connected components (two nodes belong to the same weak or strong component only if there is a path connecting them in either or both directions, respectively); the normalized in/out degree for each node (number of edges with that node as the target/source); and the normalized in/out-degree centrality (number of connections to/from the column, equal to the total weight of incoming/outgoing edges).

For the full-connectivity network, initiation zones (characterized by low mean onset delay, Figure S8 *second row*) generally correspond to lower in-degree and out-degree regions (Figure S8 *bottom row*) . For the local connectivity network, regions with high percent time active (corresponding to participating regions during Up states in Figure 5F and Figure S9, *second row*) have high in/out-degree and centrality (Figure S9, *bottom rows*) .

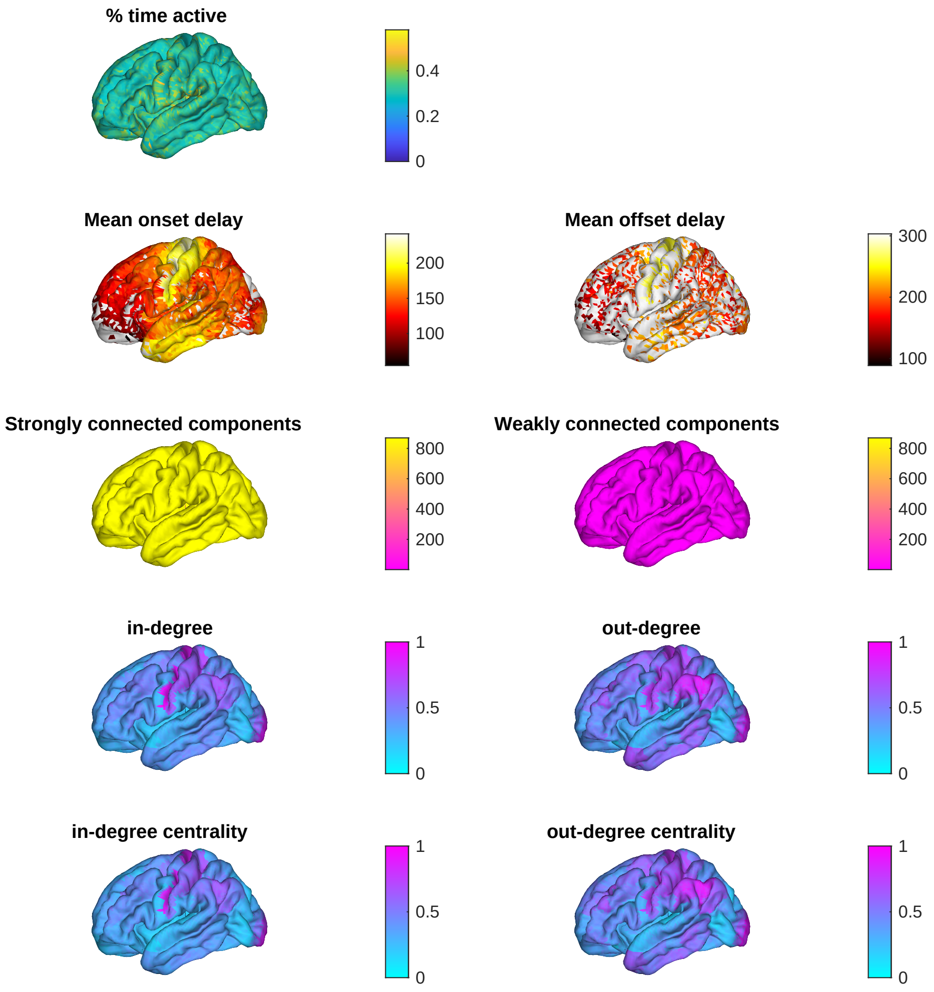


**Figure S8.** Activity maps and graph properties for the full-connectivity, Global SO network in Figure 2. Rows from top to bottom show (1) the percent of simulation time that each pyramidal cell in layer 2 spends in an Up state; (2) the mean onset/offset delay across Up states for each pyramidal cell in layer 2; (3) the number of strongly/weakly connected components in the structural connectivity graph, constructed with each cortical column as a node and each directed edge weight as the number of synaptic connections from one column to another; (4) the normalized in/out-degree for each node in the graph; and (5) the normalized in/out-degree centrality for each node in the graph, defined as the number of synaptic connections to/from that column (see Graph Properties for details). Initiation zones (low mean onset delay) at the front and back and have generally lower in-degree and out-degree than areas in the middle.

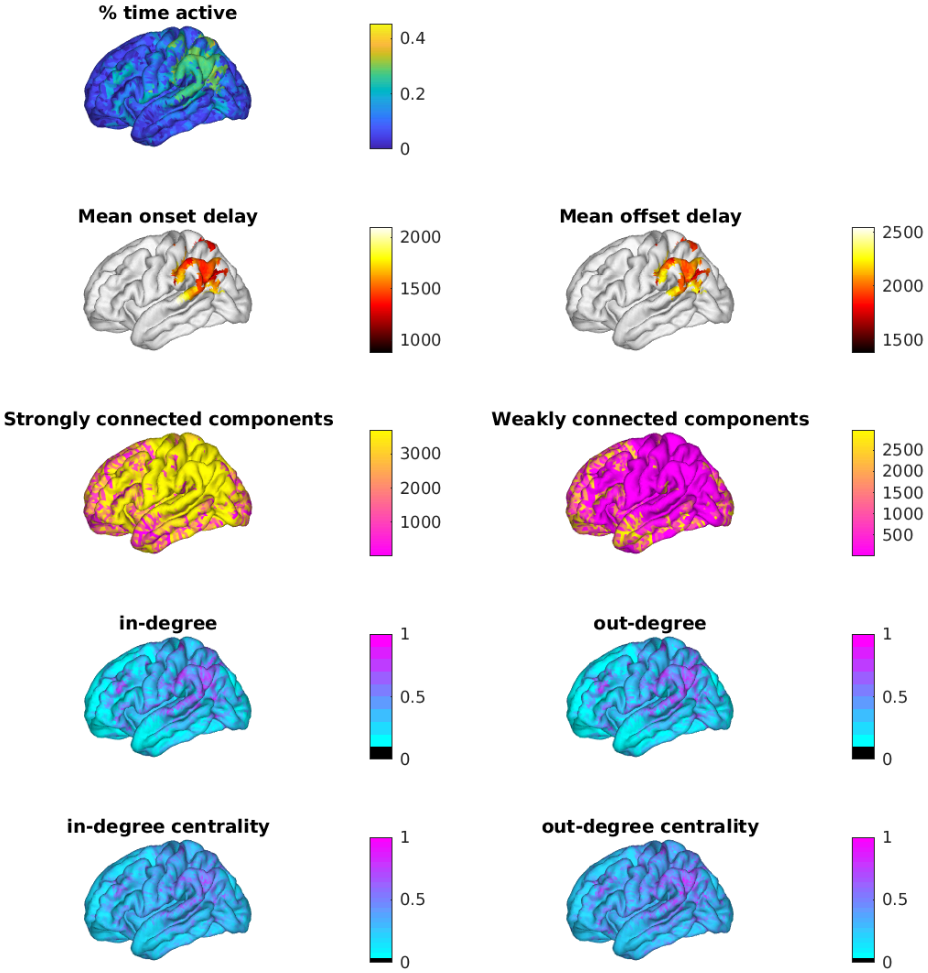


**Figure S9.** Activity maps and graph properties for the $R = 2.5$mm network in Figure 5. Rows from top to bottom show (1) the percent of simulation time that each pyramidal cell in layer 2 spends in an Up state; (2) the mean onset/offset delay across Up states for each pyramidal cell in layer 2; (3) the number of strongly/weakly connected components in the structural connectivity graph, constructed with each cortical column as a node and each directed edge weight as the number of synaptic connections from one column to another; (4) the normalized in/out-degree for each node in the graph; and (5) the normalized in/out-degree centrality for each node in the graph, defined as the number of synaptic connections to/from that column (see Graph Properties for details). Regions with high percent time active have high in/out-degree and centrality

### Model Coherence

Table S1 lists all frequencies used for coherence analysis.

| Frequency (Hz) | Bandwidth (Hz) |
|---|---|
| **0.5** | 0.5 |
| **1** | 0.5 |
| **1.5** | 0.5 |
| **2** | 0.5 |
| **2.5** | 0.5 |
| **3** | 0.5 |
| **3.5** | 0.5 |
| **4** | 0.5 |
| **4.5** | 0.5 |
| **5** | 0.5 |
| **6** | 1 |
| **7** | 1 |
| **8** | 1 |
| **9** | 1 |
| **10** | 1 |
| **12** | 2 |
| **14** | 2 |
| **16** | 2 |
| **18** | 2 |
| **20** | 2 |
| **22** | 2 |
| **24** | 2 |
| **26** | 2 |
| **30** | 2 |
| **35** | 5 |
| **40** | 5 |
| **50** | 10 |
| **60** | 10 |
| **70** | 10 |
| **80** | 10 |
| **90** | 10 |
| **100** | 10 |

**Table S1.** Peaks and 3 dB attenuation bandwidths for the second-order resonator filters used to extract narrow-band signals.

**Video S1.** The base global slow wave sleep model over 30 seconds of simulated time. The top plot shows Layer 2 cell voltages across the cortex, while the bottom plot shows the average voltage trace.

**Video S2.** The slow wave sleep model with a connection density of P = 0.1 over 30 seconds of simulated time. The top plot shows Layer 2 cell voltages across the cortex, while the bottom plot shows the average voltage trace.

**Video S3.** The slow wave sleep model with a connection range of R = 2.5 mm over 20 seconds of simulated time. The top plot shows Layer 2 cell voltages across the cortex, while the bottom plot shows the average voltage trace.

**Video S4.** The slow wave sleep model with connections longer than 10mm reduced 5-fold over 105 seconds of simulated time. The top plot shows Layer 2 cell voltages across the cortex, while the bottom plot shows the average voltage trace.
